# Diversity and genomic determinants of the microbiomes associated with COVID-19 and non-COVID respiratory diseases

**DOI:** 10.1101/2020.10.19.345702

**Authors:** M. Nazmul Hoque, M. Shaminur Rahman, Rasel Ahmed, Md. Sabbir Hossain, Md. Shahidul Islam, Keith A Crandall, Md Tofazzal Islam, M. Anwar Hossain, AMAM Zonaed Siddiki

## Abstract

The novel coronavirus disease 2019 (COVID-19) is a rapidly emerging and highly transmissible disease caused by the Severe Acute Respiratory Syndrome CoronaVirus-2 (SARS-CoV-2). Understanding the microbiomes associated with the upper respiratory tract infection (URTI), chronic obstructive pulmonary disease (COPD) and COVID-19 diseases has clinical interest. We hypothesized that the diversity of microbiome compositions and their genomic features are associated with different pathological conditions of these human respiratory tract diseases (COVID-19 and non-COVID; URTI and COPD). To test this hypothesis, we analyzed 21 whole metagenome sequences (WMS) including eleven COVID-19 (BD = 6 and China = 5), six COPD (UK = 6) and four URTI (USA = 4) samples to unravel the diversity of microbiomes, their genomic features and relevant metabolic functions. The WMS data mapped to 534 bacterial, 60 archaeal and 61 viral genomes with distinct variation in the microbiome composition across the samples (COVID-19>COPD>URTI). Notably, 94.57%, 80.0% and 24.59% bacterial, archaeal and viral genera shared between the COVID-19 and non-COVID samples, respectively, however, the COVID-19 related samples had sole association with 16 viral genera other than SARS-CoV-2. Strain-level virome profiling revealed 660 and 729 strains in COVID-19 and non-COVID sequence data, respectively and of them 34.50% strains shared between the conditions. Functional annotation of metagenomics sequences of thevCOVID-19 and non-COVID groups identified the association of several biochemical pathways related to basic metabolism (amino acid and energy), ABC transporters, membrane transport, replication and repair, clustering-based subsystems, virulence, disease and defense, adhesion, regulation of virulence, programmed cell death, and primary immunodeficiency. We also detected 30 functional gene groups/classes associated with resistance to antibiotics and toxic compounds (RATC) in both COVID-19 and non-COVID microbiomes. Furthermore, a predominant higher abundance of cobalt-zinc-cadmium resistance (CZCR) and multidrug resistance to efflux pumps (MREP) genes were detected in COVID-19 metagenome. The profiles of microbiome diversity and associated microbial genomic features found in both COVID-19 and non-COVID (COPD and URTI) samples might be helpful for developing the microbiome-based diagnostics and therapeutics for COVID-19 and non-COVID respiratory diseases. However, future studies might be carried out to explore the microbiome dynamics and the cross-talk between host and microbiomes employing larger volume of samples from different ethnic groups and geoclimatic conditions.

## Introduction

The pneumonia like respiratory illness (COVID-19) caused by a novel coronavirus, Severe Acute Respiratory Syndrome CoronaVirus-2 (SARS-CoV-2) was first reported in Wuhan city of China in December, 2019. The disease unprecedentedly spread across 216 countries and/or territories of the globe within months (He et al., 2020; Hoque et al., 2020a; Petersen et al., 2020; Rahman et al., 2020a). The World Health Organization (WHO) declared the pandemic as a public health emergency of international concern. Metagenomic RNA sequencing of COVID-19 suspected patient’s sample suggested the etiology as a new RNA virus belonging to *Coronaviridae* and later designated as SARS-CoV-2 (Hoque et al., 2020a; Lam et al., 2020). This virus can easily undergo mutation and recombination to adapt diverse environment (Islam et al., 2020), and thus survive by altering a wide host range causing constant and long-term health threats (Paules et al., 2020). Furthermore, COVID-19 has not only devastated the public health, but also posed an immense impact on human activities at the societal, economical, and geopolitical levels.

The human health services have also been adversely affected by the prevalence, recognition, and management of other non-COVID respiratory diseases including upper respiratory tract infection (URTI), asthma, and chronic obstructive pulmonary disease (COPD) (Stolz et al., 2019; Lippi and Henry, 2020). Although, SARS-CoV-2 affects different people in different ways, most common clinical features include fever, dry cough, tiredness, sore throat, diarrhea, difficulty in breathing or shortness of breath, chest pain or pressure, and loss of speech or movement (Hoque et al., 2020a; Islam et al., 2020), many of these symptoms are common in URTI and COPD. The inhaled SARS-CoV-2 virus particle likely binds to epithelial cells in the nasal cavity, starts replicating, migrates down the respiratory tract along the conducting airways, and a more robust innate immune response is triggered (Mason, 2020). Therefore, it can assume that during this propagation, migration and immune response, the microbiomes throughout the respiratory airways could be altered or changed, and some of them can further aggravate the disease process. Recent meta-analysis demonstrated that COPD is the most strongly predictive comorbidity of COVID-19 severity, and is associated with more than five-fold increased risk of COVID-19 infection (Lippi and Henry, 2020). Therefore, patients with a history of non-COVID respiratory diseases such as URTI, asthma and COPD, would be anticipated to have an increased risk of infection (Halpin et al., 2020).

The use of whole metagenome sequencing (WMS), which generates simultaneously genome reads from both prokaryotic cells and viruses, has promoted the microbiome studies effectively by recovering bacterial, archaeal, viral and phage genomes from metagenomes of ecological and clinical samples (Mathieu et al., 2018; Hoque et al., 2019; Zheng et al., 2019; Hoque et al., 2020b; Hoque et al., 2020c). However, only limited studies have been carried out using high-throughput metagenomics sequencing of the of SARS-CoV-2 infected samples (Lam et al., 2020; Wahba et al., 2020). Using this approach, the taxonomic and functional compositions of microbiomes, and the host-microbiomes interactions could be revealed in COVID-19 and non-COVID respiratory disease (URTI and COPD)-associated microbiomes.

Human health is the outcome of the complex interactions between the inhabiting microbiome and its human host (Kumpitsch et al., 2019). The compositional and functional perturbations of the microbiome can occur at different body sites, and thus, the microbiome dysbiosis can reduce the beneficial and/or commensal bacteria, and favors growth of the opportunistic pathogenic bacteria (Kumpitsch et al., 2019; Hoque et al., 2020b). During COVID-19, and non-COVID respiratory diseases (URTI and COPD), a paradigm shifts in microbiome compositions and their diversity might be observed due to the emergence of particular dominant bacteria in lungs and/or respiratory tract (Hahn et al., 2018; Mathieu et al., 2018; Wypych et al., 2019). The microbial community of the respiratory tract and lung is continually being renewed and replaced, and displays greater variations in both taxonomic composition and diversity (Mathieu et al., 2018; Wypych et al., 2019). Even though bacteria, archaea, viruses, and bacteriophages represent a significant part of the respiratory tract associated microbial communities, to get a deeper understanding of these microbes remains challenging due to the difficulties in their isolation, identification, and characterization. Viruses are essential constituents of microbial communities contributing to their homeostasis and evolution. The viral community in the human gut microbiome is dominated by bacteriophages (Dutilh et al., 2014). Phages can modulate the structure and function of a bacterial community through horizontal gene transfer (HGT), thereby altering the bacterial phenotypes including virulence, antibiotic resistance, and biofilm formation (Huddleston, 2014; Hoque et al., 2019; Zheng et al., 2019). Moreover, phage-induced alterations could pose potential health risks by influencing bacterial pathogenicity and antibiotic resistance (Zheng et al., 2019).

A large body of literature is available on characterization of the clinical features of the COVID-19 patients, the genomic features including structure, composition, evolution, and genome-wide variations in different strains of SARS-CoV-2, and to develop effective vaccine candidates and therapeutics to halt this pandemic disease (Chen et al., 2020; Hoque et al., 2020a; Huang et al., 2020; Rahman et al., 2020a; Rahman et al., 2020b; Rahman et al., 2020c). Moreover, several preliminary studies suggested that the composition of a person’s gut microbiome may be predictive of developing severe symptoms of SARS-CoV2 infections (Chen et al., 2020; He et al., 2020). However, little is known about the potential role of nasopharyngeal and lung microbiomes, and their biological mechanisms for the variable susceptibility to the respiratory diseases. To address the above unresolved question, we investigated the composition, and diversity of microbiomes associated with COVID-19, and non COVID diseases (URTI and COPD). We identified core respiratory microbiota which predicted the proteomic biomarkers of COVID-19 and non-COVID respiratory diseases using high throughput metagenomics sequencing of 21 samples. We conducted further metabolomics analysis to unravel potential biological mechanisms linking microbiome changes, SARS-CoV-2 genomic diversity, and host disease state. This reports for the first time describe the association of a diverse groups of microbiotas (bacteria, archaea and viruses) and their concomitant genomic features in COVID-19 and non-COVID (URTI and COPD) diseases, and discusses their roles the pathophysiology of these diseases.

## Methods

### Sample collection and confirmatory diagnosis of COVID 19

The nasopharyngeal specimens were collected from patients presenting with potential COVID-19 infections following the guidance of the Director General of Health Services of Bangladesh. A single dry swab was inserted through one nostril straight back (NOT upwards), along the floor of the nasal passage up to the posterior wall of the nasopharynx, and rubbed against and above the nasal turbinate. The swab was rotated a few times against the nasal wall before removing. A second swab was used to abrade the tonsils and pharynx. The swabs were placed in sample collection vial containing normal saline, and preserved at −80 °C until further use for RNA extraction. The confirmatory diagnosis of SARS-CoV-2 infections were made by RT-qPCR (Péré et al., 2020).

### RNA extraction, metagenomics library preparation and sequencing

We further intended to characterize the microbial community present along with the SARS-CoV-2 virus in the nasopharyngeal specimens of COVID-19 patients through whole metagenome sequencing (WMS). We utilized total RNA-Seq approach for metagenomics study, and total RNA was extracted from five (n=5) COVID-19 positive patients using TRIzol (Invitrogen) reagent following manufacturer’s protocol. RNA-Seq libraries were prepared from isolated total RNA using TruSeq Stranded Total RNA Library Prep kit (Illumina) according to the manufacturer’s instructions where first strand cDNA was synthesized using SuperScript II Reverse Transcriptase (Thermo Fisher), and random primers (Visnovska et al., 2019; Zhou et al., 2020). Paired-end (2 x 150 bp reads) sequencing of the RNA library was performed on the Illumina NextSeq 500 platform. RNA extraction, library preparation, and sequencing were carried out at the Basic and Applied Research on Jute Project, Dhaka, Bangladesh.

### Sequence retrieval

In addition to our five COVID-19 metagenome sequences, we retrieved six (n=6) Chinese COVID-19 metagenome sequences from the NCBI (National Center for Biotechnology Information) database (https://www.ncbi.nlm.nih.gov/) (Accession numbers: SRX7705831-SRX7705836), four (n=4) shotgun metagenome sequences of human URTI belonged to CDC, USA from the NCBI (Accession numbers: SRR10252885, SRR10252888, SRR10252889 and SRR10252892 under bio-project: PRJNA573045), and six (n=6) metagenome sequences of human COPD from the European Nucleotide Archive, UK (Accession numbers: ERR2732537, ERR2732541, ERR2732559, ERR2732558, ERR2732551 and ERR2732550 under bio-project: PRJEB14074). Therefore, the current metagenome study included 21 sequences including 11 COVID-19 (BD=5, China=6) and 10 non-COVID (USA=4, UK=6) sequences to investigate the microbiome diversity and composition, and metabolic functional potentials associated with COVID 19 (BD and China), and non-COVID respiratory diseases (COPD; UK and URTI; USA).

### Taxonomic mapping, classification, diversity and community analysis

The shotgun WMS data were analyzed using the assembly-based hybrid method MG-RAST (release version 4.1) (Glass et al., 2010). The paired-end FASTQ files were concatenated and filtered through BBDuk (Stewart et al., 2018) (with options k = 21, mink = 6, ktrim = r, ftm = 5, qtrim = rl, trimq = 20, minlen = 30, overwrite = true) to remove Illumina adapters, known Illumina artifacts, and phiX. Any sequence below these thresholds or reads containing more than one ‘N’ were discarded. In the current study, 46.04 million reads in two metagenomes with an average of 2.19 million reads per sample passed the quality control steps (Supplementary Data 1). The trimmed sequences were uploaded to the MG-RAST server with properly embedded metadata and were subjected to optional quality filtering with dereplication and host DNA removal, and finally screening for taxonomic and functional assignment. Alpha diversity (diversity within samples) was estimated using the observed species, and Shannon diversity indices (Koh, 2018) MG-RAST read assignments and counts. To visualize differences in bacterial diversity, principal coordinate analysis (PCoA; at genus level) was performed based on the Bray-Curtis distance method (Beck et al., 2013). Taxonomic abundance was determined by applying the ‘‘Best Hit Classification” option using the NCBI database as a reference with the following settings: maximum *e*-value of 1×10^-30^; minimum identity of 80% for bacteria, 60% for archaea and viruses, and a minimum alignment length of 20 as the set parameters. A ‘target’ genome library was constructed containing all viral sequences from the NCBI RefSeq Release 201 database (https://en.wikipedia.org/wiki/National_Center_for_Biotechnology_Information) using the Kraken 2 (Wood et al., 2019), and the metagenomics reads were then aligned against the target library using the BWA algorithm (Jaillard et al., 2016). Finally, the PathoScope 2 was used for strain level taxonomic assignment of the viruses (Hong et al., 2014). In all of the bioinformatics analysis, the Chinese sequences were used only as forward sequences.

### Functional profiling of the microbiomes

We performed the taxonomic functional classification through mapping the reads onto the Kyoto Encyclopedia of Genes and Genomes (KEGG) database (Kanehisa et al., 2019), and SEED subsystem identifiers (Glass et al., 2010) on the MG-RAST server using the partially modified set parameters (*e-*value cutoff: 1×10^-30^, min. % identity cutoff: 60%, and min. alignment length cutoff: 20).

### Statistical analysis

The non-parametric test Kruskal-Wallis rank sum test was used to evaluate differences in the relative percent abundance of taxa in COVID-19 and non-COVID (URTI and COPD) sequence data. Comparative taxonomic and functional profiling was performed with the reference prokaryotic metagenomes available in MG-RAST database for statistical analyses. The gene counts were normalized by dividing the number of gene hits to individual taxa/function by total number of gene hits in each metagenome to remove bias due to differences in sequencing efforts. To identify differentially abundant SEED or KEGG functions, and resistance to antibiotics and toxic compounds (RATCs) across the four sampling locations (BD, China, UK and USA), statistical tests were applied with non-parametric test Kruskal-Wallis rank sum test at different KEGG and SEED subsystems levels IBM SPSS (SPSS, Version 23.0, IBM Corp., NY, USA).

## Results

### Microbiome diversity and compositions in COVID-19 and non-COVID samples

The COVID-19 disease significantly affected the diversity (Shannon estimated alpha diversity, P=0.036, Kruskal-Wallis test) and composition (Bray-Curtis distance estimated beta diversity, P=0.021, Kruskal-Wallis test) of respiratory tract microbiome (Fig. 1). The rarefaction curves showed that sequencing depth was sufficient to capture the entire microbial diversity among the samples of the study (Fig. S1). Both the number of identified microbial taxa and Shannon estimated alpha diversity were significantly higher in COVID-19 samples (BD and China) than the non-COVID (UK and USA) samples (Fig. 1A). The principal coordinate analysis based on Bray-Curtis distances showed distinct separations among the four sampling locations (Bangladesh, China, USA, and UK) (Fig. 1B). Notably, the COVID-19 related samples (BD and China) remained more similar than the URTI and COPD-associated samples from the USA and UK (Fig. 1B). Moreover, across the non-COVID respiratory diseases (URTI and COPD)-associated samples of UK and USA, all of the samples clustered more closely than those of COVID-19 samples (BD and China) indicating that SARS-CoV-2 infections are associated with diverse secondary microbial communities (Fig. 1B).

**Fig. 1.**
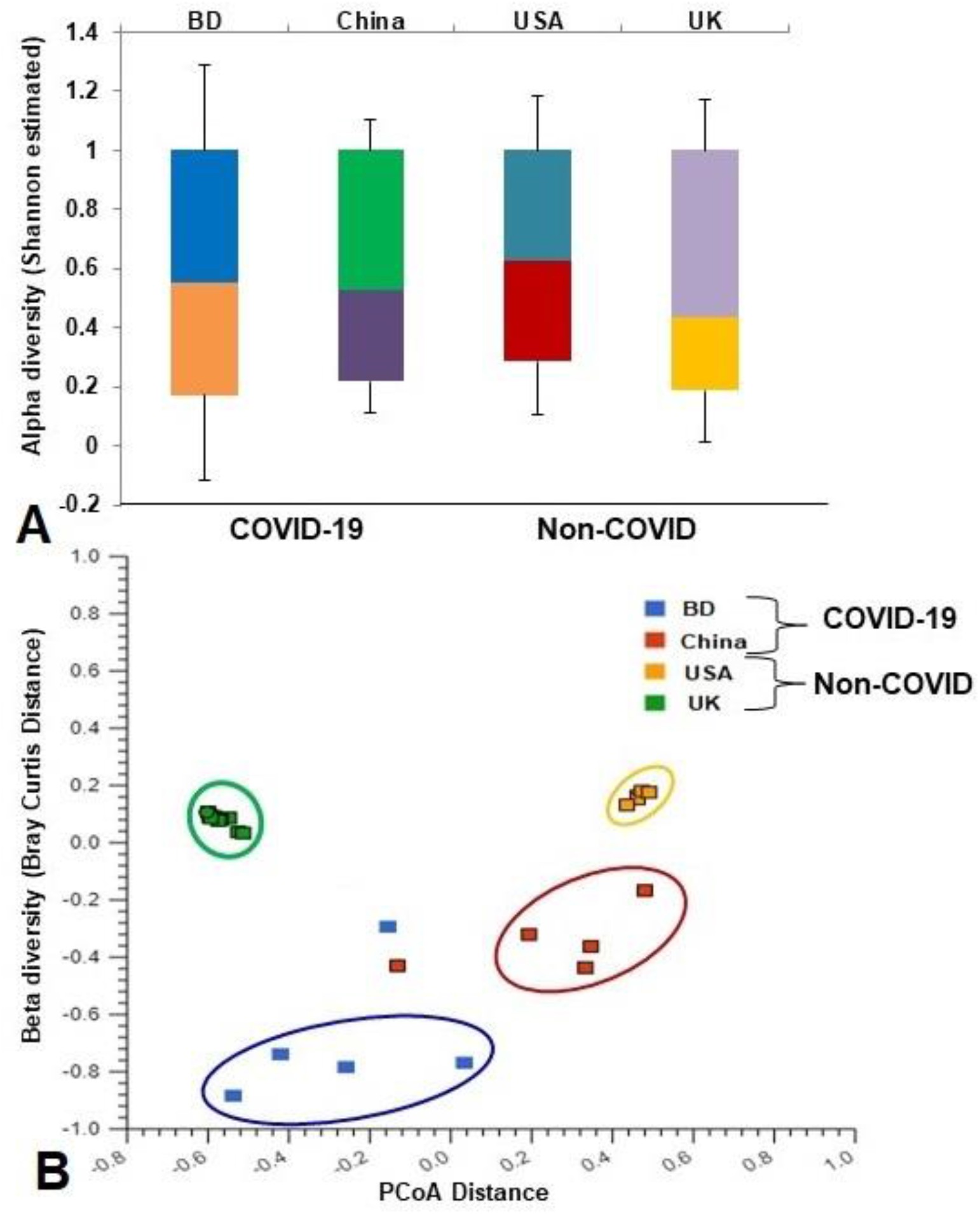
Differences in microbiome diversity and community structure in COVID-19 (BD and China), and non-COVID (UK and USA) disease metagenomes. (A) Box plots showing significant differences (P=0.036, Kruskal-Wallis test) in Shannon estimated alpha diversity in four metagenomes. (B) Principal coordinates analysis (PCoA) measured on the Bray-Curtis distance method separated samples by microbial population structure. Each dot represents an individual, and colors indicate the populations in four metagenomes. Statistical analysis using Kruskal-Wallis tests showed significant microbial diversity variations across the four metagenomes (P=0.021, Kruskal-Wallis test).

The composition of COVID-19 (BD and China) and non-COVID respiratory disease (URTI; USA and COPD; UK)-related microbiomes at the domain level was dominated by bacteria, with a relative abundance of >99.50, followed by archaea (~ 0.3%), and viruses (~0.2%) (Data S1). Twenty-eight bacterial phyla were identified in COVID-19 (BD and China), and non-COVID (UK and USA) metagenomes including 28, 10, 28 and 27 phyla in BD, China, UK and USA samples, respectively (Data S1). A total of 534 bacterial genera were detected across the four sample locations including 527, 306, 512 and 443 in BD, China, UK and USA samples, respectively (Fig. 2A, Data S1). About 95% (505/534) of the detected bacterial genera shared in both COVID-19 and non-COVID diseases, and only 22 genera had unique association with COVID-19 disease (Fig. 2 A, Data S1). In addition to bacteria, 60 archaeal and 61 viral genera were detected, and of them 48, 10, 60 and 31 archaeal genera (Fig. S2B), and 31, 5, 36 and 18 viral genera (Fig. S2C) were identified in BD, China, UK and USA, respectively. Of the detected archaeal genera, majority of the genera (80.0%) were found to be shared in both sample groups, and only 20.0% (12/60) archaeal genera had unique association with non-COVID diseases (Fig. 2B, Data S1). Similarly, 26.23% and 49.18% of the detected viral genera had unique association with COVID-19 and non-COVID diseases, respectively, and 24.59% genera were found to be shared between the conditions (Fig. 2C, Data S1). The strain level profiling of the viral communities revealed 1032 strains including 660 and 729 in the COVID-19 and non-COVID samples, respectively (Fig. 2 D, Data S1). Of the identified strains, 303 in COVID-19 and 373 in non-COVID respiratory disease samples, respectively had unique associations, and 34.50% strains were found to be shared between the conditions (Fig. 2 D, Data S1).

**Fig. 2.**
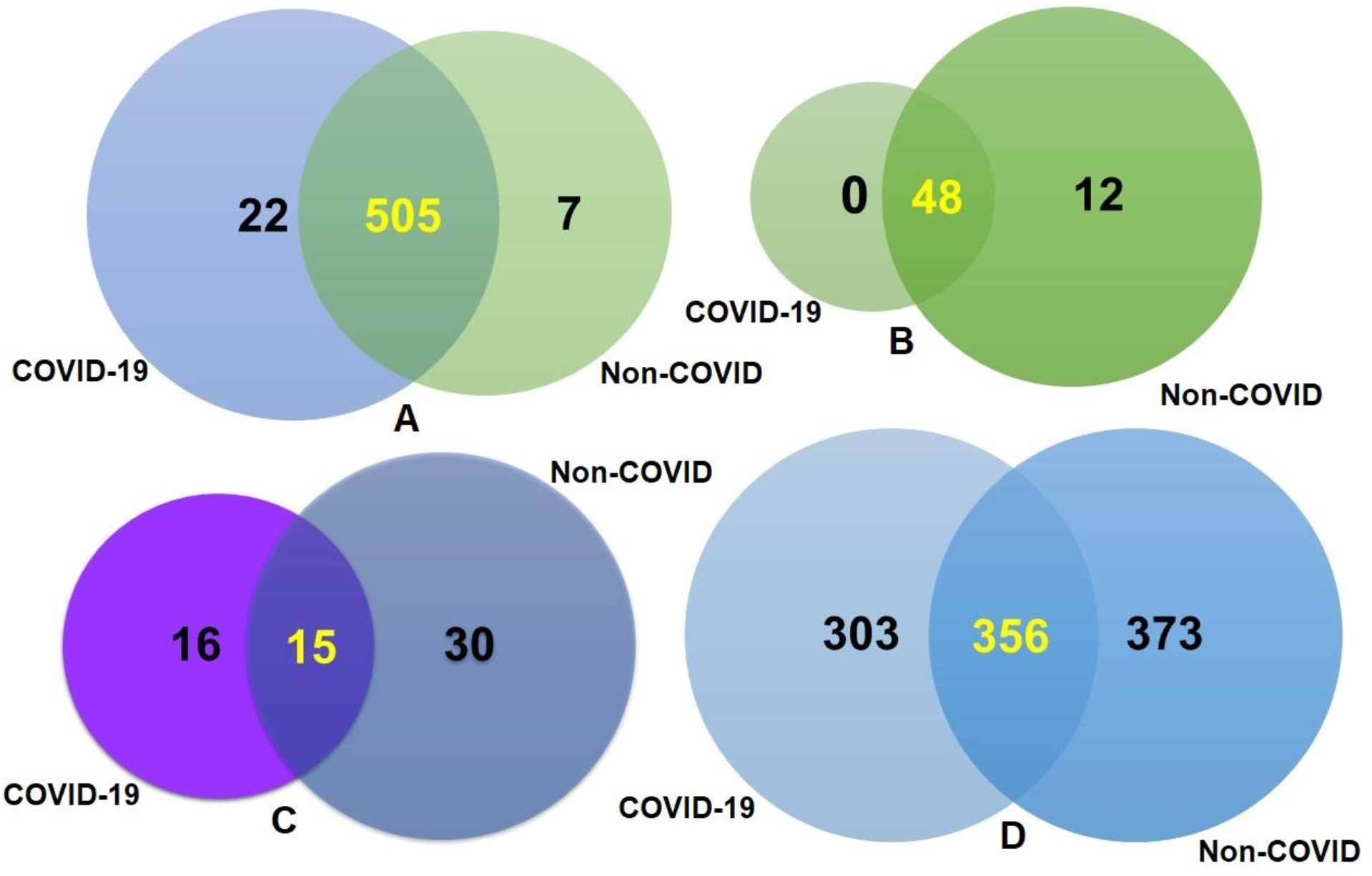
Taxonomic composition of COVID-19 (BD and China) and non-COVID (URTI; USA and COPD; UK) disease metagenomes. Venn diagrams representing the core unique and shared microbiomes in COVID-19 and non-COVID diseases. (A) Venn diagram showing unique and shared bacterial genera. Out of 534 detected bacterial genera, only 13 genera had unique association with COVID-19 while rest of the 521 genera (highlighted in yellow) shared between the condition. (B) Venn diagram comparison of unique and shared archaeal genera where 48 (highlighted in yellow) genera shared between the condition, and 12 genera had unique association with non-COVID URTI and COPD) diseases. (C) Venn diagrams representing unique and shared viral genera identified in both metagenomes. Of the detected viral genera (n=61), 16 and 30 genera had unique association with COVID-19 and non-COVID diseases, respectively, and 15 genera (highlighted in yellow) were found to be shared between the study metagenomes. (D) Venn diagrams showing the unique and shared viral strains in COVID-19 and Non-COVID diseases. Out of viral strains detected, 356 strains (highlighted in yellow) shared between the condition while 303 and 373 strains had unique associations with COVID-19 and Non-COVID metagenomes, respectively. More information on the taxonomic results are available in Data S1.

### Differences in bacteriome compositions in COVID-19 and non-COVID samples

The composition of bacterial phyla across the two metagenome groups varied significantly (P=0.037, Kruskal-Wallis test). This diversity was represented by the predominant bacterial phyla of *Proteobacteria* (35.59%), *Tenericutes* (18.09%), *Actinobacteria* (17.42%), *Cyanobacteria* (11.23%), *Firmicutes* (7.60%), and *Bacteroidetes* (6.20%) in the COVID-19 metagenome, and *Firmicutes* (56.47%), *Bacteroidetes* (14.59%), *Actinobacteria* (14.12%), and *Fusobacteria* (2.38%) in non-COVID samples (Fig. 3; Data S1). The rest of the phyla in four geographic locations had relatively lower abundances (< 1.0%) (Data S1).

The composition and relative abundance of the bacteria significantly differed (P = 0.029, Kruskal-Wallis test) at the genus level between COVID-19 (BD and China) and non-COVID (UK and URTI) disease samples. Remarkably, most of the genera had higher relative abundances in the COVID-19 samples compared to the non-COVID samples (Data S1). Importantly, *Staphylococcus* (10.39%), *Nostoc* (6.70%), *Anabaena* (4.86%), *Mycobacterium* (3.02%), *Cyanothece* (2.51%), *Bradyrhizobium* (2.32%), *Actinomyces* (1.81%), *Pseudomonas* (1.80%), *Propionibacterium* (1.56%), *Corynebacterium* (1.52%), *Rhodopseudomonas* (1.30%), *Nodularia* (1.27%), *Burkholderia* (1.25%), *Micrococcus* (1.23%), *Acinetobacter* (1.18%), *Methylobacterium* (1.15%), *Streptomyces* (1.14%), *Rhodococcus* (1.09%), and *Rhodobacter* (1.00%) were the most abundant genera in COVID-19 samples (Fig. 4, Data S1). Conversely, *Prevotella* (18.20%), *Streptococcus* (17.35%), *Veillonella* (12.64%), *Rothia* (9.70%), *Actinomyces* (7.97%), *Neisseria* (2.55%), *Fusobacterium* (2.13%), *Lactobacillus* (1.97%), *Atopobium* (1.53%), *Megasphaera* (1.45%), *Porphyromonas* (1.41%), *Bacteroides* (1.34%), *Peptostreptococcus* (1.28%), *Clostridium* (1.24%), and *Bifidobacterium* (1.05%) were the predominantly abundant genera in non-COVID (COPD and URTI) samples of the UK and USA (Fig. 4, Data S1). The rest of the genera detected in both COVID-19 and non-COVID respiratory disease group had less than 1.00% relative abundances (Data S1).

**Fig. 3.**
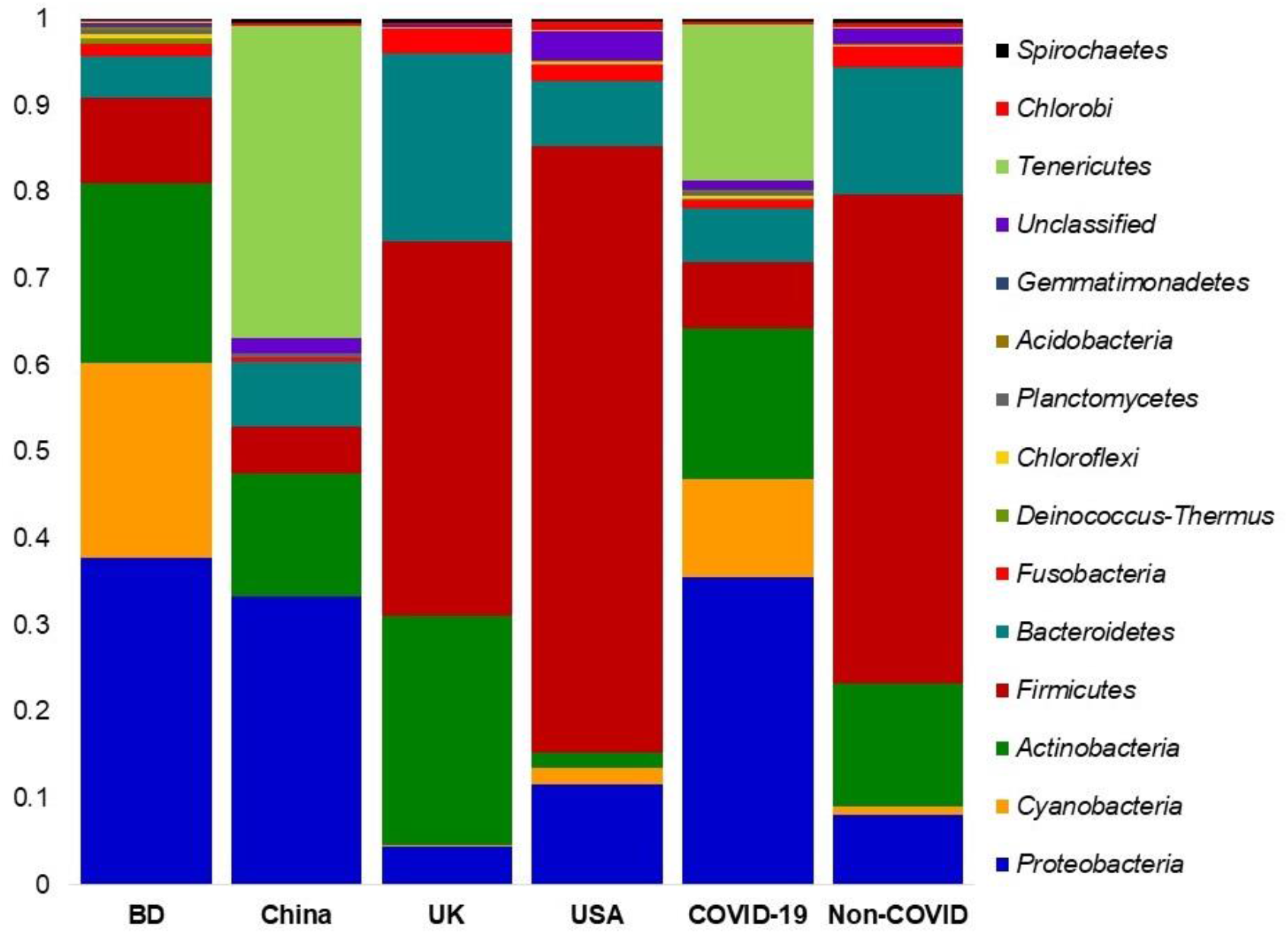
The phylum level taxonomic profile bacteria in in COVID-19 (BD and China), and respiratory tract (UK and USA) disease metagenomes. Stacked bar plots showing the relative abundance and distribution of the 15 top abundant phyla, with ranks ordered from bottom to top by their increasing proportion. The first four bar plots represent the abundance of bacterial phyla in the corresponding category of COVID-19 (BD and China) and non-COVID (URTI; USA and COPD; UK) disease, and the last two bar plots depict overall relative abundance of phyla in COVID-19 and non-COVID metagenomes, respectively. The distribution and relative abundance of the bacterial phyla in the study metagenomes are also available in Data S1.

**Fig. 4.**
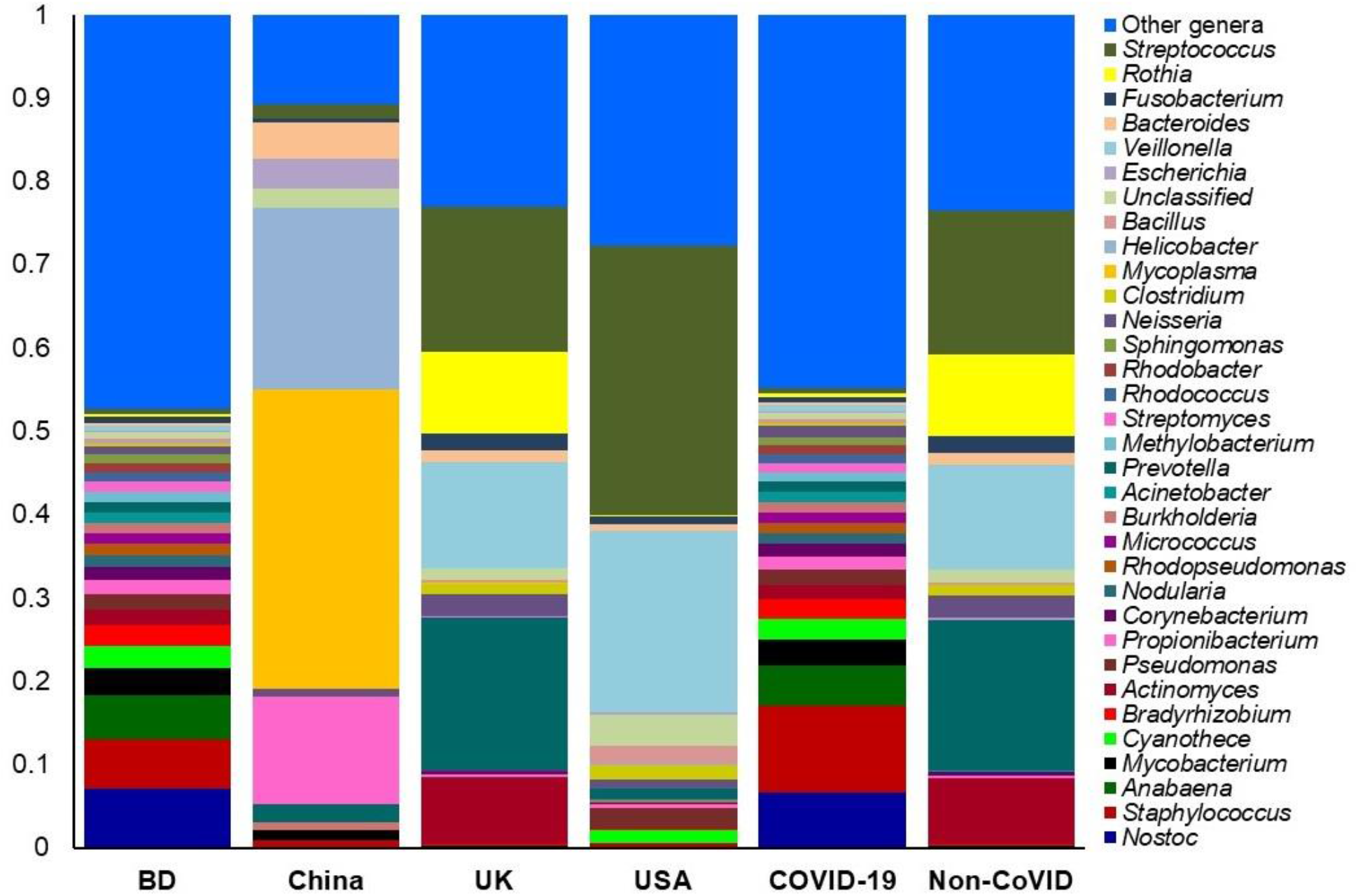
The genus level taxonomic profile bacteria in COVID-19 (BD and China) and non-COVID (URTI; USA and COPD; UK) disease metagenomes. Stacked bar plots showing the relative abundance and distribution of the 34 most top abundant bacterial genera, with ranks ordered from bottom to top by their increasing proportion. The first four bar plots represent the abundance of bacteria in the corresponding category of COVID-19 (BD and China) and non-COVID (URTI; USA and COPD; UK) disease, and the last two bar plots depict overall relative abundance of bacterial genera in COVID-19 and non-COVID metagenomes, respectively. Only the 33 most abundant genera are shown in the legend, with the remaining genera grouped as ‘Other genera’. The distribution and relative abundance of the bacterial genera in the study metagenomes are also available in Data S1.

### Differences in archaea and viruses in COVID-19 and non-COVID samples

The archaeal and viral fractions of microbiomes in COVID-19 (BD and China), and non-COVID respiratory disease (COPD; UK and URTI; USA)-samples were concomitantly detected. The composition and relative abundance of these two domains varied significantly (P = 0.017, Kruskal-Wallis test) between the COVID-19 and non-COVID samples (Data S1). Among the identified archaeal components of the microbiome, *Methanosarcina* (19.3%), *Methanocaldococcus* (17.76%), *Thermococcus* (10.30%), *Methanothermobacter* (8.28%), *Haloarcula* (7.77%), *Staphylothermus* (6.01%), *Natronomonas* (5.38%), *Ferroglobus* (3.52%), *Caldivirga* (3.38%), *Halobacterium* (2.83%), *Natrialba* (2.18%), *Methanosphaerula* (2.08% and *Picrophilus* (2.02%) were the predominantly abundant genera in COVID-19 samples (Fig. 5, Data S1). Conversely, *Methanobrevibacter* (10.84%), *Methanococcus* (10.17%), *Methanocorpusculum* (6.97%), *Pyrococcus* (4.76%), *Methanosphaera* (4.24%), *Methanococcoides* (3.52%), *Methanosaeta* (2.55%), *Archaeoglobus* (2.48%), *Methanospirillum* (2.40%) and *Methanoculleus* (2.26%) were the most abundant archaeal genera in the non-COVID respiratory disease associated samples (Fig. 5, Data S1). Though the rest of the archaeal genera had lower relative abundances (< 2.0%) across the samples of both categories, however, their abundances always remained higher in COVID-19 related BD and non-COVID associated UK (COPD) samples (Fig. 5, Data S1).

**Fig. 5.**
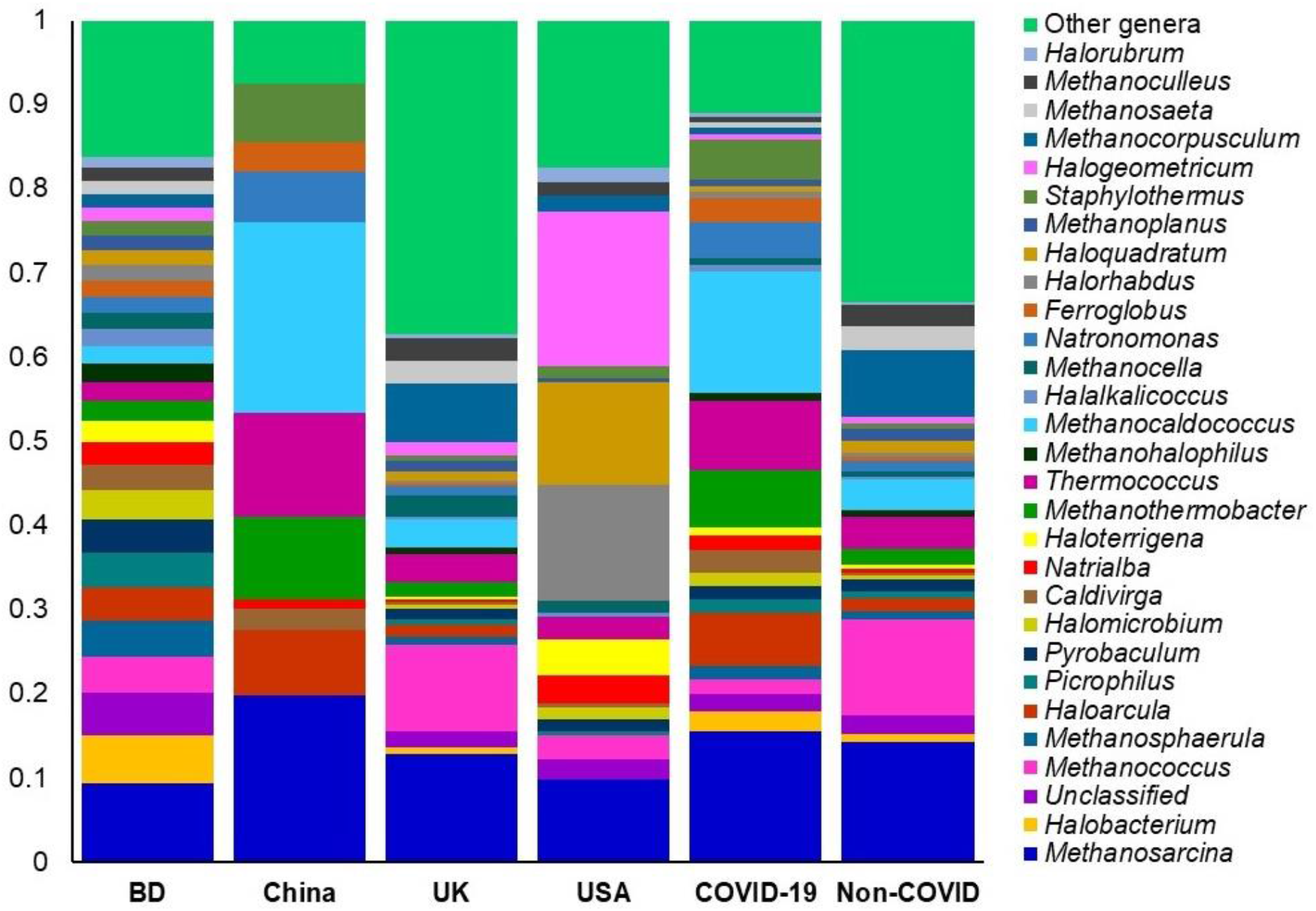
The genus level taxonomic profile archaea in COVID-19 (BD and China) and respiratory tract disease (UK and USA) metagenomes. Stacked bar plots showing the relative abundance and distribution of the 30 most abundant genera, with ranks ordered from bottom to top by their increasing proportion. The first four bar plots represent the abundance of archaea in the corresponding category of COVID-19 (BD and China) and non-COVID (URTI; USA and COPD; UK) disease, and the last two bar plots depict overall relative abundance of archaeal genera in COVID-19 and non-COVID metagenomes, respectively. Only the 29 most abundant genera are shown in the legend, with the remaining genera grouped as ‘Other genera’. The distribution and relative abundance of the archaeal genera in the study metagenomes are also available in Data S1.

The viral fraction of the microbiome was largely dominated by *Betacoronavirus* (99.67% of the total abundance) genus in the COVID-19 related samples (BD; 99.55% and China; 99.98%) while the relative abundance of this genus was 39.78% in the non-COVID samples. In addition, *Siphovirus* (33.17%), *Alphapapillomavirus* (12.01%) and *Myovirus* (7.16%) were the most predominant viral genera in non-COVID samples (Fig. 6, Data S1). Rest of the viral genera detected across the both sample groups had lower relative abundances (<1.0%) (Data S1). The COVID-19 samples from BD and China had sole association of 16 viral genera (other than *betacoronavirus*), and of them, *Tombusvirus*, *Victorivirus*, *Partitivirus*, *Chrysovirus* and *Totivirus* were the most abundant genera (Data S2).

**Fig. 6.**
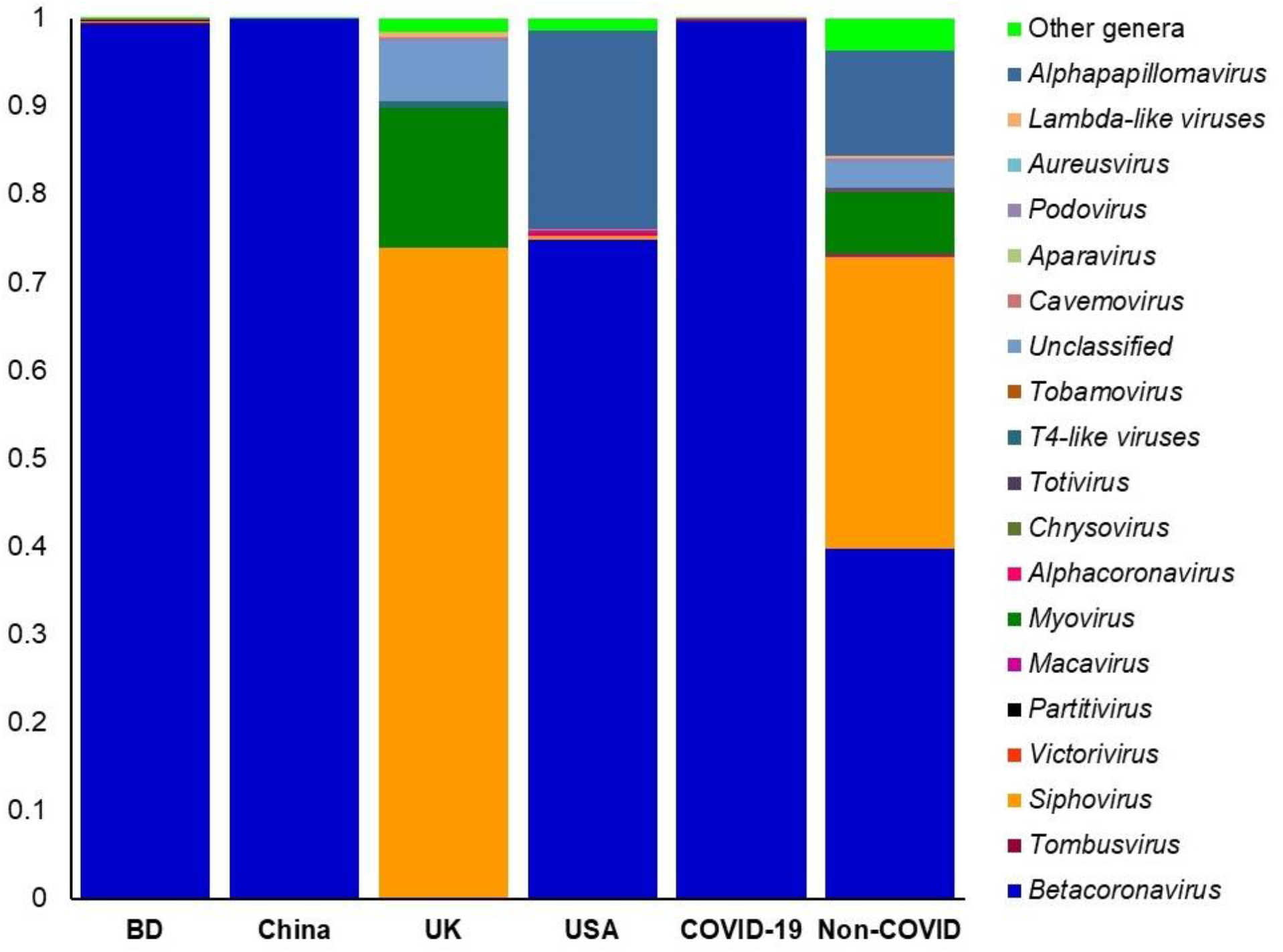
The taxonomic profile virus in in COVID-19 (BD and China) and respiratory tract disease (UK and USA) metagenomes. Stacked bar plots showing the relative abundance and distribution of the 20 viral genera, with ranks ordered from bottom to top by their increasing proportion among the BD, China, UK and USA metagenomes. The first four bar plots represent the abundance of viruses in the corresponding category of COVID-19 (BD and China) and non-COVID (URTI; USA and COPD; UK) disease, and the last two bar plots depict overall relative abundance of viral genera in COVID-19 and non-COVID metagenomes, respectively. Only the 19 most abundant genera are shown in the legend, with the remaining genera grouped as ‘Other genera’. The distribution and relative abundance of the viral genera in the study metagenomes are also available in Data S1.

Complex microbial communities can harbor multiple strains of the same species which was evident in the present study since the strains of associated viral genera varied significantly (P = 0.033, Kruskal-Wallis test) between the COVID-19 and non-COVID samples. The relative abundances of the identified viral strains also varied between the sample categories since the COVID-19 related samples had the single most predominantly abundant (63.78%) strain of SARS-CoV-2, and this strain was not detected in the non-COVID samples. The COVID-19 related samples of BD and China also possessed *Choristoneura fumiferana* granulovirus (13.17%), *Shamonda orthobunyavirus* (3.64%) and *Tupaiid betaherpesvirus* 1 (1.46%) (Fig. 7A, Data S1). On the other hand, *Simbu orthobunyavirus* (13.26%), *Proteus* phage VB_PmiS-Isfahan (11.37%), *Mycobacterium* phage Enkosi (9.22), *BeAn* 58058 virus (8.27%), *Kaisodi* virus (5.00%), *Mycobacterium* phage Phrux (3.33%), *Beilong* virus (2.88%), *Mollivirus sibericum* (2.77%), *Oxbow virus* (2.21%) and *Micromonas* sp. RCC1109 virus MpV1 (2.1%) were the top abundant viral strains in non-COVID samples of UK and USA (Fig. 7B, Data S1). Phylogenetic analysis revealed that the SARS-CoV-2 strain found in COVID-19 patients showed neighboring relationship to *SARS Coronavirus* Tor2, *Beihai hermit* crab virus 4 and *Penicillium aurantiogriseum* partitivirus 1 (Fig. 7A).

**Fig. 7:**
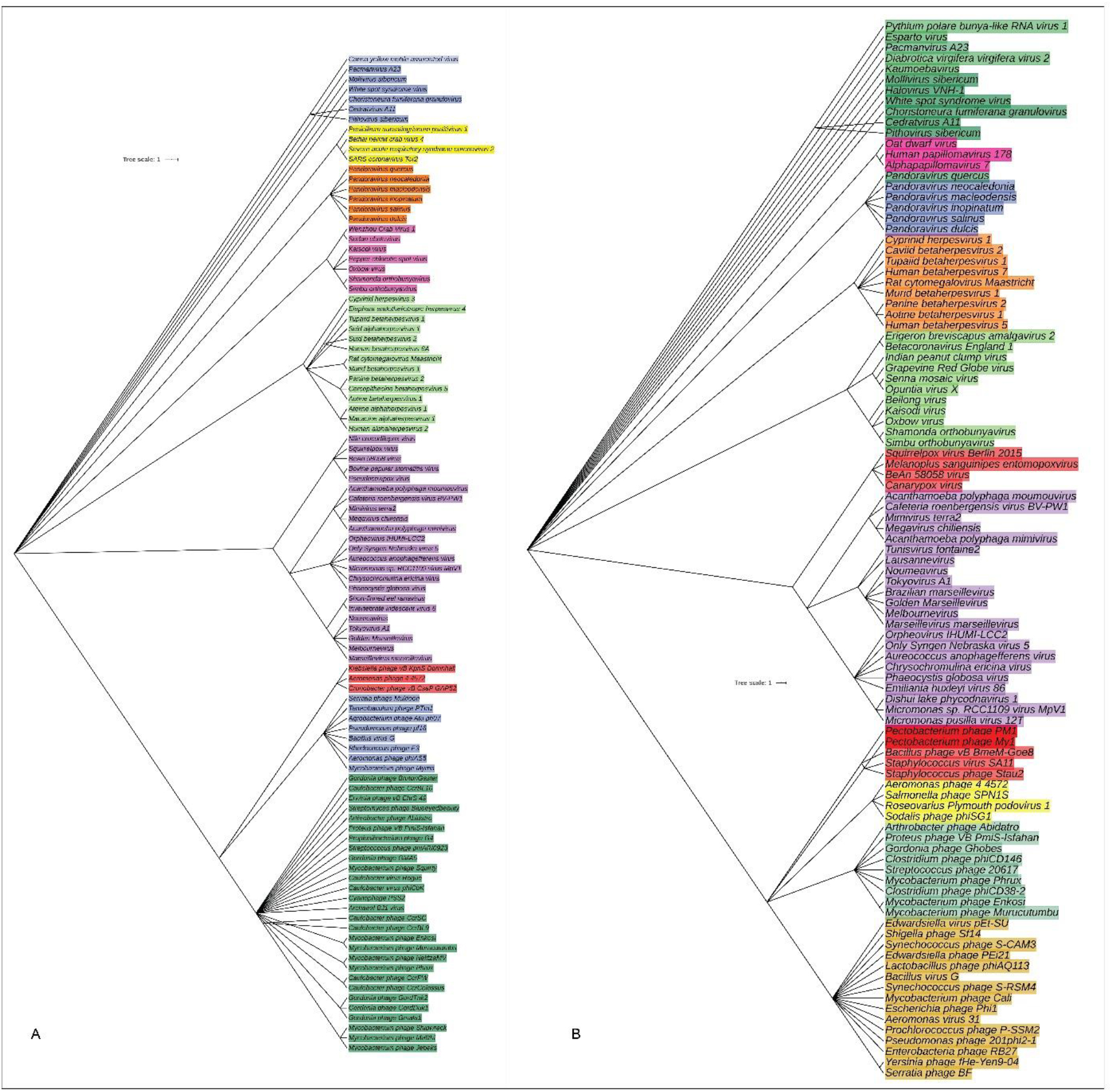
The species and/or strain level taxonomic representation of viruses in COVID-19 and non-COVID (URTI and COPD) metagenomes. Sequences are assigned to different taxonomic index in PathoScope (PS) analysis using minimum identity of 95% and minimum alignment length 20 as cutoff parameters. The slanted phylogenetic trees were generated with the top 100 abundant strains of viruses in the COVID-19 (A) and non-COVID (B) metagenomes based on the maximum likelihood method using the NCBI taxonomy tree and visualized with iTOL (interactive Tree Of Life). The bootstrap considered 1000 replicates. The scale bar represents the expected number of substitutions averaged over all the analyzed sites. The length of the scale bar represents 1 nucleotide substitution per 100 positions. Different colors are assigned according to the taxonomic ranks of the viruses. The species and/or strains used in the phylogenetic tree are also available in Data S1.

### Differences in metabolic functions in COVID-19 and non-COVID samples

By comparing the number of genes assigned to each KEGG pathway among the groups, we found a series of significant differences (P=0.006, Kruskal-Wallis test) that lead to the functional divergence in the COVID-19 (BD and China), and non-COVID respiratory disease (COPD; UK and URTI; USA)-associated microbiomes. In the comparative analysis of predicted KEGG orthology (KO), we found that the COVID-19 associated microbiomes harbored relatively higher abundance of genes coding for amino acid metabolism (55.23%; BD=63.79% and China=76.67%), energy metabolism (11.01%; BD=9.81% and China=12.22%), membrane transport (68.00%; BD=62.09% and China=73.90%), ABC transporters (67.89%; BD=89.91% and China=65.87%), replication and repair (28.74%; BD=23.25% and China=34.23%), flagellar assembly (42.22%; BD=35.31% and China=49.13%), and primary immunodeficiency diseases (56.18%; BD=71.11% and China=63.24%) compared to the non-COVID related microbiomes of COPD (UK) and URTI (USA)-associated metagenomes (Fig. 8A, Data S2). On the other hand, the non-COVID respiratory disease (COPD and URTI)-associated microbiomes had over expression of genes coding for oxidative phosphorylation (61.30%; UK=67.22% and USA=55.38%), signal transduction (50.91%; UK=46.08% and USA=75.74%), infectious diseases (59.95%; UK=67.11% and USA=52.79%), bacterial chemotaxis (53.79%; UK=85.00% and USA=22.58%), and bacterial secretion systems (29.38%; UK=23.76% and USA=35.00%) than the COVID-19 related microbiomes (Fig. 8A, Data S2).

**Fig. 8.**
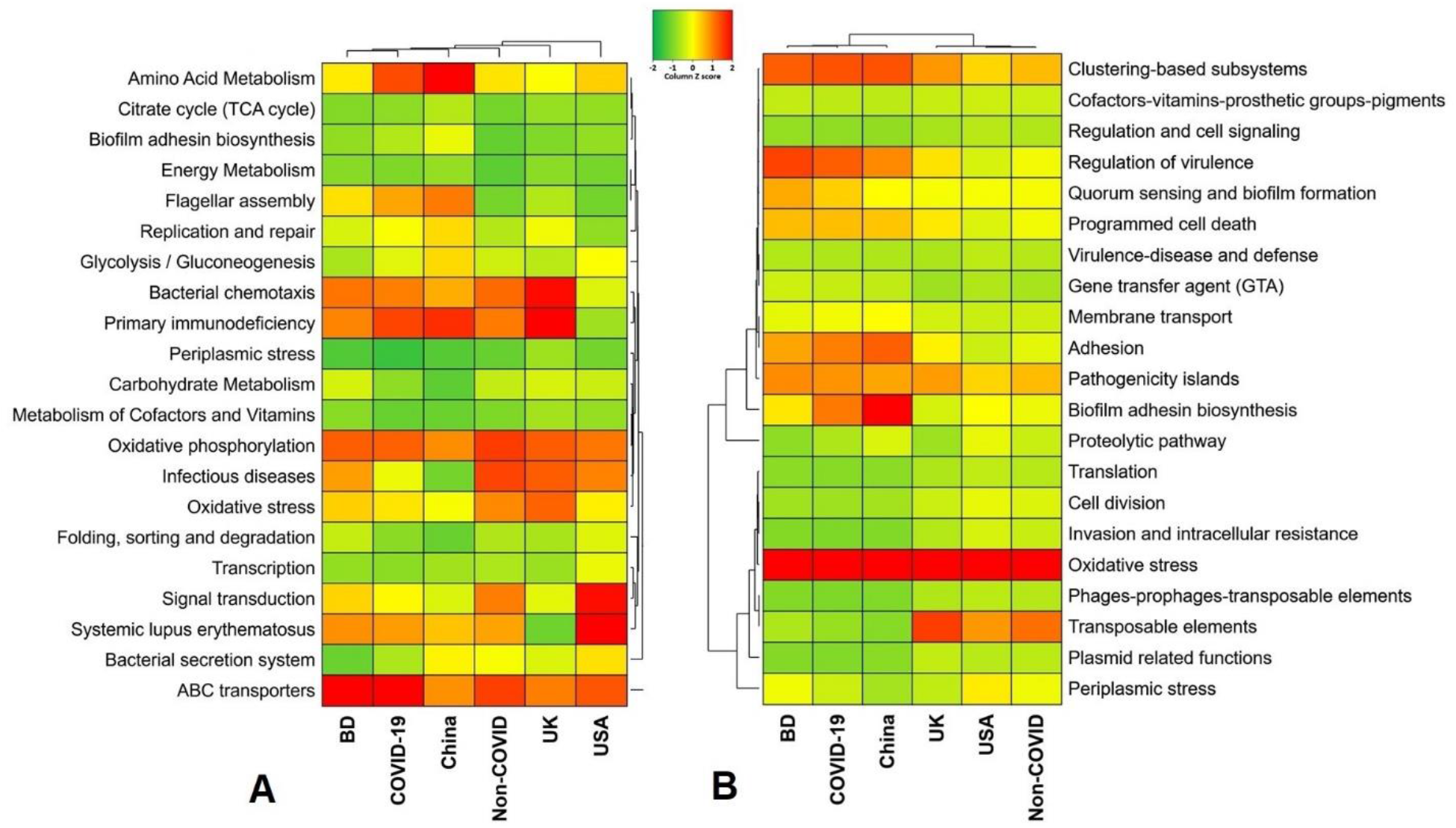
Functional annotation of the COVID-19 (BD and China) and respiratory tract disease (UK and USA) metagenomes. (A) Heatmap representing the average relative abundance hierarchical clustering of the predicted KEGG Orthologs (KOs) functional pathways of the microbiome across four metagenome groups. (B) Heatmap showing the average relative abundance hierarchical clustering of the predicted SEED functions in different levels among the microbiomes of four metagenomes. The color bars (column Z score) at the top represent the relative abundance of putative genes. The color codes indicate the presence and completeness of each KEGG and SEED module, expressed as a value between −2 (lowest abundance) and 2 (highest abundance). The red color indicates the more abundant patterns, whilst green cells account for less abundant putative genes in that particular metagenome.

The SEED subsystems in the COVID-19 (BD and China) and non-COVID (COPD; UK and URTI; USA)-related microbial communities significantly (P=0.024, Kruskal-Wallis test) varied. The COVID-19 related microbiomes had relatively higher abundance of genes encoding for pathogenicity islands (21.55%; BD=18.69% and China=24.42%), clustering-based subsystems (21.15%; BD=22.23% and China=20.07%), regulation of virulence (20.34%; BD=24.41% and China=16.28%), biofilm adhesion biosynthesis (18.49%; BD=11.58% and China=25.40), adhesion (17.96%; BD=16.77% and China=19.13%), programmed cell death (13.53%; BD=14.85% and China=12.21%), membrane transport (6.66%; BD=7.83% and China=8.50%), gene transfer agent (5.00%; BD=5.88% and China=4.12%) and virulence, disease and defense (3.39%; BD=3.78% and China=3.00%) (Fig. 8B, Data S2). In contrast, genes coding for oxidative stress (48.75%; UK=41.89%, USA=65.60%), periplasmic stress (6.49%; UK=10.40%, USA=2.56), cell division (4.65%; UK=3.29%, USA=6.01%), invasion and intracellular resistance (2.88%; UK=1.70%, USA=4.06%), and phages, prophages and transposable elements (1.26%; UK=1.40%, USA=1.11%) were upregulated in the non-COVID related COPD and URTI causing microbiomes (Fig. 8B, Data S2).

### Differences in resistomes composition in microbiomes of COVID-19 and non-COVID samples

The resistome analysis of the COVID-19 (BD and China) and non-COVID (UK and USA)-associated microbiomes, the SEED module of the MG-RAST pipeline provided a comprehensive scenario. Using SEED, 150,146 reads mapped to 30 resistance to antibiotics and toxic compounds (RATC) functional groups across the sample across the four geographic regions, with different relative abundances (Data S2). The RATC genes were classified into two unique groups such as antibiotic resistance (n=17) and toxic metal resistance (n=13) group. Among the RATC functional groups, cobalt-zinc-cadmium resistance (CZCR) had higher relative abundances in COVID-19 microbiomes (23.69%; BD=25.85%, China=21.53%) compared to non-COVID (10.34%; UK=11.75%, USA=8.92%) microbiomes (Fig. 9, Data S2). Moreover, the RATC functional genes associated with beta lactamase (*BLAC*) (13.55%; BD=9.23% and China=17.87%), copper homeostasis (*CH*) (11.18%; BD=12.33% and China=10.02%), multidrug resistance to efflux pumps (*MREP*) (10.97%; BD=14.75% and China=7.19%), multidrug resistance cluster (*mdt*ABCD) (5.34%; BD=4.57% and China=6.11%), arsenic resistance (*AR*) (4.76%; BD=6.27% and China=3.26%) and multidrug-efflux-pump in *Campylobacter jejuni* (*Cme*ABC) operon (1.86%; BD=2.74% and China=0.98%) had higher relative abundances in the COVID-19 associated microbiomes than the non-COVID respiratory disease (COPD and URTI) causing microbiomes (Fig. 9; Data S2). Conversely, gens coding for resistance to fluoroquinolones (*RFL*) (22.31%; UK=26.16%, USA=18.46%), *Bla*R1-Family-Regulatory-Sensor-Transducer-Disambiguation (*Bla*R1) family (15.47%; UK=9.71% and USA=21.23%), methicillin resistance in *Staphylococci (MRS)* (8.20%; UK=10.55% and USA=5.85%), resistance to chromium compounds (*RCHC*) (5.11%; UK=1.05%, USA=9.23%), *Streptococcus pneumonia-vancomycin-* tolerance-locus (*SPVANT*) (3.39%; UK=3.71%, USA=3.07%), mercury resistance to operon (*MRO*) (2.34%; UK=2.53% and USA=2.15%) and copper homeostasis: copper tolerance (*CHCT*) (2.07%; UK=2.91%, USA=1.23%) were found to be over expressed in the COPD and URTI-associated non-COVID microbiomes (Fig. 9, Data S2). Rest of the RATC genes detected in both metagenomes also varied in their relative abundances.

**Fig. 9.**
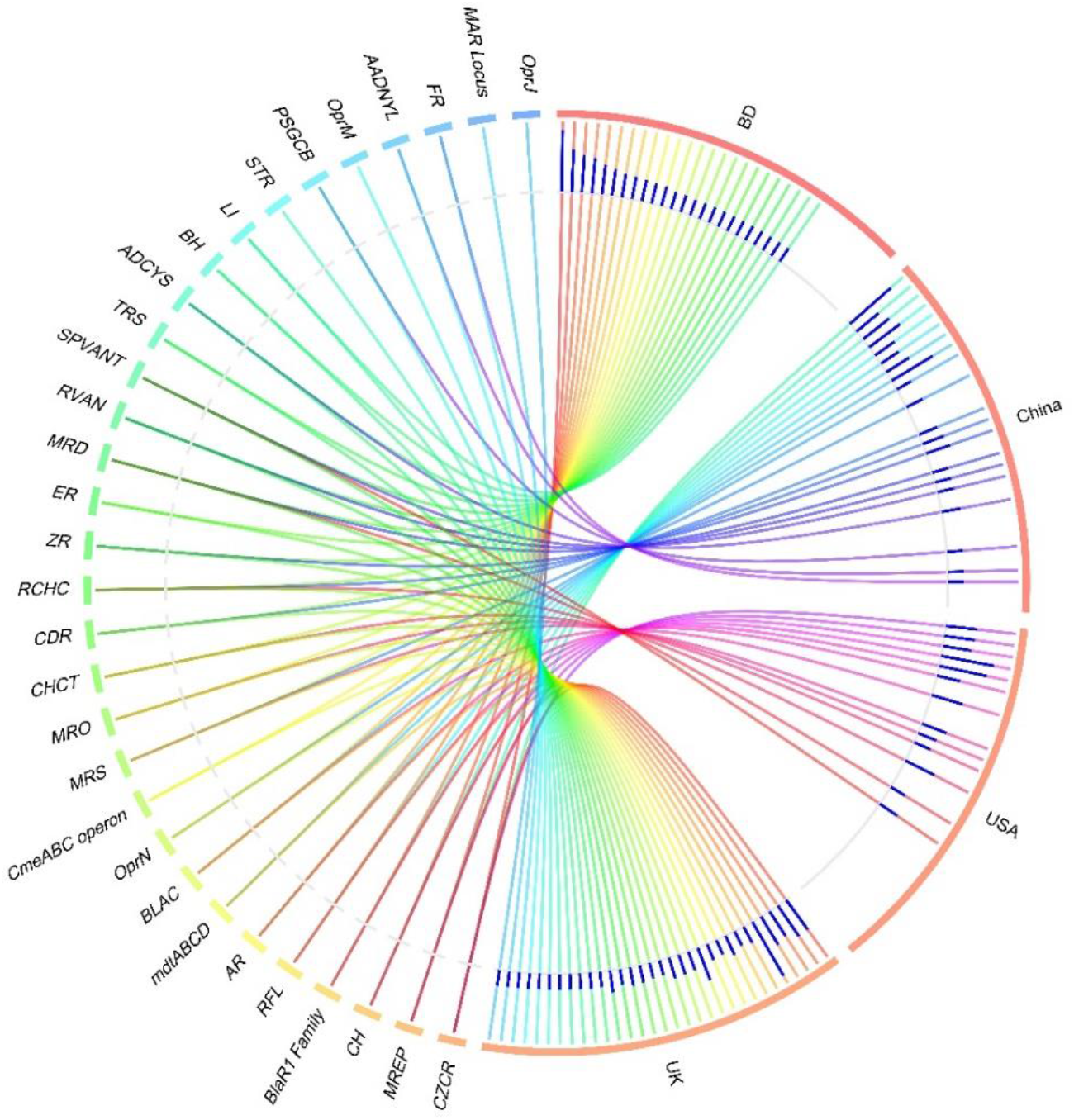
Distribution of the resistance to antibiotic and toxic compounds (RATC) genes in COVID-19 (BD and China) and respiratory tract disease (UK and USA) metagenomes. The circular plot illustrates the diversity and relative abundance of the RATC genes detected among the microbiomes of the four metagenomes through SEED subsystems analysis. The association of the RATC genes according to metagenome is shown by different colored ribbons and the relative abundances these genes are represented by inner blue colored bars. Part of the RATC functional groups are shared among microbes of the four metagenomes (BD, China, UK and USA), and some are effectively undetected in the microbiomes of the other metagenomes. Abbreviations-*CZCR*: cobalt-zinc-cadmium resistance, *MREP*: multidrug resistance to efflux pumps, *CH*: copper homeostasis; *BlaR1* Family: BlaR1 family regulatory sensor-transducer disambiguation, *RFL*: resistance to fluoroquinolones, *AR*: arsenic resistance, *mdtABCD*: the mdtABCD multidrug resistance cluster, *BLAC*: beta-lactamase resistance, *OprN*: mexe-mexf-oprn multidrug efflux system, *CmeABC* operon: multidrug efflux pump in *Campylobacter jejuni, MRS*: methicillin resistance in *Staphylococci, MRO*: mercury resistance to operon, *CHCT*: copper homeostasis: copper tolerance, *CDR*: cadmium resistance, *RCHC*: resistance to chromium compounds, *ZR*: zinc resistance, *ER*: erythromycin resistance, *MRD*: mercuric reductase, *RVAN*: resistance to vancomycin, *SPVANT: Streptococcus pneumonia* vancomycin tolerance locus, *TRS*: teicoplanin-resistance in *Staphylococcus, ADCYS*: adaptation to d-cysteine, *BH*: bile hydrolysis, *LI*: lysozyme inhibitors, *PSGCB*: polymyxin synthetase gene cluster in *Bacillus, OprM*: mexA-mexB-oprm multidrug efflux system, *AADNYL*: aminoglycoside adenylyltransferases, *FR*: fosfomycin resistance, *MAR* Locus: multiple antibiotic resistance to locus, *OprJ*: mexC-mexD-OprJ-multi drug-efflux-system.

## Discussion

The present study represents a proof-of-concept to decipher changes in microbiome compositions and diversity in COVID-19 (BD and China), and non-COVID related chronic obstructive pulmonary disease; COPD (UK), and upper respiratory tract infection; URTI (USA). Several recent studies showed that COVID-19 is associated with dysbiosis of the gut microbiomes (Dhar and Mohanty, 2020; Zuo et al., 2020). Respiratory tract especially lung microbiome is more dynamic and transient than that of the gastrointestinal tract because of bidirectional movement of air and mucus (Huffnagle et al., 2017). This makes it plausible that SARS-CoV-2 infections could lead to abnormal inflammatory reactions to worsen the symptoms of COVID-19. The microbiome diversity (alpha and beta diversity) measures revealed higher microbial diversity in COVID-19 samples which remained more similar than the URTI and COPD-associated samples from the USA and UK. Regardless of higher taxonomic abundances, the COVID-19-associated microbiomes remained inconsistent and fluctuates more within BD and China metagenomes than those of UK and USA samples. In a recent study, Zou et al. (2020) reported that loss of salutary species in COVID-19 persisted in most patients despite clearance of SARS-CoV-2 virus, and is associated with a more long-lasting detrimental effect to the gut microbiome.

In the present study, although the dominant microbial phyla showed no significant difference between two groups, COVID-19 still affected the prevalence of some microbes belonging to *Proteobacteria* and *Cyanobacteria* corroborating the recent findings of Khatiwada and Subedi (2020). To be specifically, in COVID-19 samples of BD and China, the abundance of *Proteobacteria*, *Tenericutes*, and *Cyanobacteria* were significantly higher. Conversely, *Firmicutes*, *Actinobacteria*, *Bacteroidetes*, and *Fusobacteria* were the predominant phyla in COPD and URTI samples of UK and USA, respectively. These findings are accorded with previous studies of respiratory tract disease associated microbiome dysbiosis (Singh et al., 2017; Hahn et al., 2018; Li et al., 2019). The healthy respiratory tract and lungs microbiomes are mostly represented by *Firmicutes, Bacteroidetes* and *Proteobacteria* phyla (Khatiwada and Subedi, 2020) suggesting inclusion of other predominant opportunistic phyla such as *Tenericutes, Cyanobacteria, Actinobacteria,* and *Fusobacteria* in the COVID-19 metagenomes of BD and China. The COVID-19 and non-COVID (COPD and URTI)-patients had an enrichment of pathogenic and commensal bacteria, indicating a degree of microbial changes in diseased states. We demonstrated that in the SARS-CoV-2 infected patients, some genera including *Staphylococcus*, *Nostoc*, *Anabaena*, *Mycobacterium*, *Cyanothece*, *Bradyrhizobium*, *Actinomyces*, *Pseudomonas*, *Propionibacterium*, *Corynebacterium*, *Rhodopseudomonas*, *Nodularia*, *Burkholderia*, *Micrococcus*, *Acinetobacter*, *Methylobacterium*, *Streptomyces*, *Rhodococcus*, and *Rhodobacter* had higher relative abundances compared to the non-COVID samples. The changes in the composition and relative abundances of the microbiomes in the respiratory tract is associated with many vital body functions such as immune regulation and pathogenesis (Kumpitsch et al., 2019), and might be associated with the COVID-19 exacerbations.

The pathophysiology of an inflammation plays an important role in destroying the invading microbes, and thereby protecting the commensal organisms (Hoque et al., 2020b), but unregulated inflammation is an underlying cause of many chronic diseases like COPD. Our findings demonstrated that the non-COVID related COPD and URTI patients had the higher relative abundance of certain dominant bacterial genera including *Prevotella*, *Streptococcus*, *Veillonella*, *Rothia*, *Actinomyces*, *Neisseria, Fusobacterium*, *Lactobacillus*, *Atopobium*, *Megasphaera*, *Porphyromonas*, *Bacteroides*, *Peptostreptococcus*, *Clostridium*, and *Bifidobacterium*. The inflammatory mediators are associated with parts of dominant microbiome in in the lungs, which suggesting that inflammation might impair the structure of growth environment of microbiome and the microbiome changes might in turn promoted the COPD (Ditz et al., 2020) and URTI (Man et al., 2017) inflammations. The upper respiratory tract is generally considered to be a major reservoir for potential pathogens, including *Streptococcus, Veillonella, Pseudomonas, Clostridium, Rothia*, and *Neisseria*, to expand and subsequently spread towards the lungs, which could potentially lead to a chronic infection like COPD (Man et al., 2017; Ditz et al., 2020). The findings of the present study suggested that bacteriome composition in COVID-19 and non-COVID associated COPD and URTI might be similar at phylum level, however, could vary significantly at genus and strain levels which might be related to the age, gender, weight, food habit, and different living environment (Wang et al., 2018). Thus, establishing and maintaining a balanced microbiota in the COVID-19 and non-COVID respiratory disease (URTI and COPD) samples that is resilient to pathogenic expansion and invasion could prove vital for respiratory health. Moreover, 94.57% bacterial genera including *Staphylococcus, Mycobacterium, Burkholderia, Prevotella, Streptomyces, Neisseria, Acidovorax, Fusobacterium, Streptococcus, Erythrobacter*, and *Bacteroides* shared between COVID-19 and non-COVID (COPD and URTI) metagenomics data suggesting their diverse role in the pathophysiology of human diseases (Man et al., 2017; Ditz et al., 2020; Khatiwada and Subedi, 2020). However, the mechanisms for the pathophysiology of COVID-19 and other respiratory tract diseases (non-COVID) as well as specific host-microbiome interactions supporting the changes in microbiome compositions, are considered below and not well-established.

Viral infections predispose patients to secondary bacterial and archaeal infections, which often have a more severe clinical course (Brundage, 2006; Yildiz et al., 2018; Shen et al., 2020; Zuo et al., 2020). Until now, most of the respiratory tract disease associated microbiomes reported so far are limited to bacteriome composition (Man et al., 2017; Singh et al., 2017; Li et al., 2019; Ditz et al., 2020; Shen et al., 2020) while a plethora of other concomitant microbial components including archaea and viruses could also be detected (Koskinen et al., 2017; Hoque et al., 2019; Hoque et al., 2020b). Unlike bacteria, the diversity and composition of archaea and viruses always remained much lower in COVID-19 and non-COVID (URTI and COPD) samples. The COVID-19 and non-COVID related COPD samples had higher number of archaeal and viral genera with significantly varied relative abundances. Among the identified archaeal components of the microbiomes, the methanogenic archaea *Methanosarcina* was the most abundant genus in both sample groups. Most methanogenic archaea coexist and interact closely with anaerobic bacteria (Hoque et al., 2020b). Methanogenic archaea utilize low molecular weight compounds, such as H2 + CO2, formic acid, or acetate, and therefore have symbiotic relationships with the producers of these substrates. Therefore, it is reasonable to hypothesize that the presence or increase in level of methanogenic archaea alters the composition of the polymicrobial community thus resulting in changes in virulence of the flora (Maeda et al., 2013; Hoque et al., 2020b). The viral fraction of the microbiome was largely dominated by *Betacoronavirus* genus in the COVID-19 related samples of BD and China. Conversely, the virome fraction of the non-COVID (URTI and COPD) samples were mostly represented by the genera belonged to the order *Caudovirales* which consists of the three families of tailed bacterial viruses (bacteriophages) infecting bacteria and archaea (Hoque et al., 2019; Hoque et al., 2020b). The host range of *Caudovirales* is very broad and includes all major bacterial phyla found in our samples: *Firmicutes, Bacteroidetes, Proteobacteria* and *Actinobacteria.* This corresponded with an increased relative abundance of these bacterial taxa in COPD and URTI metagenomes. Moreover, the presence of few predominating viral genera both in COVID-19 and non-COVID samples suggests that the crucial differences might be occurring at the strain level, and most of the genus identified in each sample were represented by a single strain. The COVID-19 related samples were dominated by a strain of betacoronavirus, the Severe Acute Respiratory Syndrome Coronavirus 2 (SARS-CoV-2) (not detected in non-COVID samples). Notably, the other predominant viral species/strains in COVID-19 and non-COVID samples were mostly represented by bacteriophages including *Proteus* phage VB_PmiS-Isfahan, *Mycobacterium* phage Enkosi, *Mycobacterium* phage Phrux. Bacteriophages are naturally occurring viruses that use bacteria as hosts, and play an extremely important part in allowing relatively harmless bacteria to become pathogens (Huddleston, 2014; Hoque et al., 2019; Zheng et al., 2019). Recent scientific researches provide evidence that bacteriophages are overlooked human pathogens, implied in the triggering, and worsening of a number of human diseases (Zheng et al., 2019). The COVID-19 causing SARS-CoV-2 strain showed neighboring relationship to human classic coronavirus, the *SARS Coronavirus* isolate Tor2 (SARS-CoV Tor2) corroborating with the recent findings of Hou (2020) and Konno et al. (2020) (Hou, 2020; Konno et al., 2020). However, our current findings demonstrated that archaea and viruses (other than SARS-CoV-2) neither cause COPD and URTI directly nor play a role in the initiation of the disease process, but later, when bacterial infection of the respiratory tract occurs, they replicate in the immune and epithelial cells, and may act as a predisposing factor as well as a primary etiological agent for more severe URTI, and prolonged disease like COPD.

The functional metabolome analyses provided a promising opportunity for studying microbial-host interactions, and has uncovered significant differences in microbial metabolic functions in COVID-19 (BD and China) and non-COVID (COPD; UK and URTI; USA) samples. In this study, the COVID-19 associated microbiomes harbored relatively higher abundance of genes coding for metabolism (amino acid and energy), ABC transporters, membrane transport, replication and repair, clustering-based subsystems, virulence, disease and defense, adhesion, regulation of virulence, programmed cell death, and primary immunodeficiency diseases compared to non-COVID microbiomes. However, the specific role of these metabolic pathways in the pathophysiology of the COVID-19 disease is not clearly explored still now. The metabolic activities related to virulence and immunosuppression have profound impact on SARS-CoV-2 infections as evidenced earlier from the SARS and MERS outbreaks (Thng et al., 2020). Most recently, investigators comprehensively queried the impact of the microbiome on the metabolome throughout the body, and found that that the microbiota affected the metabolomics of every organ (Hoque et al., 2020b; Lee-Sarwar et al., 2020; Quinn et al., 2020). The observed increased abundance of genes associated with primary immune diseases (e.g., adenosine deaminase) in COVID-19 and COPD pathogens is responsible for inhibition of T cell maturation and lymphocytic proliferation, very low CD4 count, cell-to-cell communication and therefore could be used as a selective marker for COVID-19 and COPD diagnosis (Hoque et al., 2019). We identified several genes as being differentially expressed in COVID-19 microbiomes that were involved in bacterial physiology, such as the bacterial secretion system by which pathogenic bacteria to secrete virulence factors for host invasion (Depluverez et al., 2016). ABC transporters, which contribute to substrate transport across the bacterial membrane and are related to antibiotic resistance (Wilson, 2016; Zhang et al., 2020), were also enriched with differentially expressed genes in the COVID-19 microbiomes.

The abundance and diversity of the antibiotic and toxic compounds (RATC) genes in the human respiratory microbiome remain poorly characterized. The microbiomes (especially bacterial community) within this potential reservoir are becoming more resistant. In the context of SARS-CoV-2 virus infection, interactions between the virus, the host, and resident bacteria with pathogenic potential are known to complicate and worsen disease, resulting in coinfection and increased morbidity and mortality of infected individuals. Recent study on antimicrobial resistance and host-microbiome interaction during influenza virus infection revealed that antimicrobial resistance plays important roles in the pathogenesis of influenza-associated bacterial secondary infection (Zhang et al., 2020) supporting our present findings. Although we may not have a complete record of past antibiotic use because antibiotics taken over-the-counter are often not reported, it is also possible that some of the RATC genes overexpressed in COVID-19 samples are unrelated to antibiotic use by the individuals in which these were detected and could be the result of transmission of antibiotic resistant strains between individuals. Transmission of multidrug resistant bacteria both within the human population and between environmental reservoirs and humans is known to occur (Bengtsson-Palme et al., 2018). Some of the RATC functional groups identified across the four sample locations are simply associated with multidrug resistant efflux pumps that naturally function to extrude toxins and chemicals from the bacteria (Alvarez-Ortega et al., 2013; Zhang et al., 2020), and thus antimicrobial resistance is a secondary effect during viral infections like SARS-CoV-2. However, the pathogenesis of SARS-CoV-2 infection and its interaction with the host metabolism should be investigated in a broader aspect to fully elucidate the role of metabolic functional potentials in COVID-19.

## Conclusions

Our metagenomics analyses revealed that COVID-19 had significant effect on the diversity and composition of respiratory tract microbiomes of human. Furthermore, the COVID-19 associated microbiomes are rich in diversity, composition and functional capacity compared to the non-COVID respiratory disease (COPD and URTI)-related microbial communities. The identifiable changes in the microbiome diversity, composition and associated genomic features demonstrated in this study might be associated with the development, treatment, and resolution of COVID-19 disease. It also highlights the importance of understanding these associations in individuals infected with SARS-CoV-2 virus and possibly other respiratory viruses. The shared bacterial, archaeal and viral genera detected in this study suggest that respiratory tract disease associated microbiomes could have a profound impact on the pathogenesis of SARS-CoV-2 infection and severity of the disease. Moreover, the predominantly abundant bacterial pathogens along with the archaea and bacterial viruses (phages) could have important contribution for the generation of immune responses against viral attack, and this secondary microbial community might affect the outcome of COVID-19 disease. The correlations of the microbiome composition and functional metabolomics in the progression of COVID-19 and non-COVID respiratory disease (COPD and URTI) determined in this study are also evidenced by the differences in the relative abundances of the genes associated with resistomes (RATC) and several metabolic functional pathways among them. Future studies should investigate cross-talks between host and respiratory tract microbiomes in a large cohort of COVID-19 patients, and correlate with the severity of this disease. Furthermore, such studies would also be enhanced by the inclusion of gut microbiomes sampling in addition to the COVID-19 sampling for direct testing of microbial transfer across the gut-lung axis.

## Data availability

The sequence data reported in this paper are available in the GISAID (a global science initiative and primary source for genomic data of influenza viruses and the novel coronavirus responsible for COVID-19) database (https://www.gisaid.org) (Accession numbers: SRX7705831-5836, SRX7705834-37), NCBI (National Center for Biotechnology Information) database (Accession number: PRJNA573045), and the ENA (European Nucleotide Archive) (Accession number: PRJEB14074).

## Ethics Statement

The protocol for sample collection from COVID-19 patients, sample processing, transport, and RNA extraction was approved by the Director General of Health Services of Bangladesh.

## Author contributions

MNH conceived and designed the study, performed bioinformatics analysis, visualized figures, interpreted results and drafted the manuscript. MSR the retrieved sequences, curated the data and performed bioinformatics analysis. RA, SH and SI performed sequencing and reviewed the drafted manuscript. MTI, KAC, MAH and AMAMZS conceived and guided the study, and critically reviewed the drafted manuscript.

## Acknowledgements

We are thankful to those who deposited and shred their sequences to the GISAID, NCBI and ENA, and make open for the global research community.

## Supplementary Information

Supplementary information supporting the findings of the study are available in this article as Data S1 and S2, and Figure S1.

