## Supplementary figures and images for "Diversity and genomic determinants of the microbiomes associated with COVID-19 and non-COVID respiratory diseases"

### Fig. S1

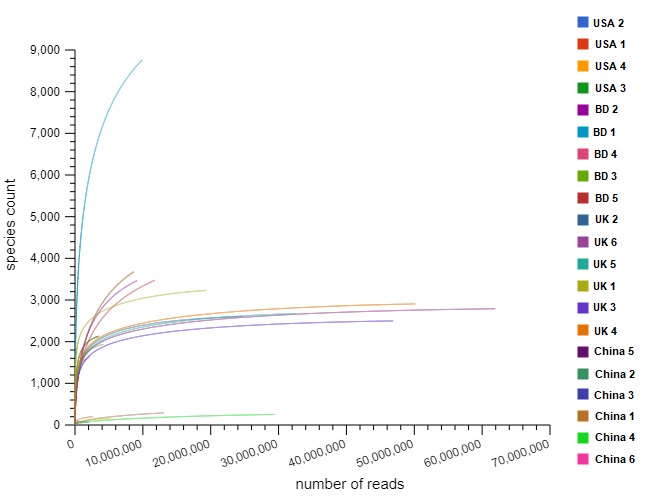

### Fig. S2

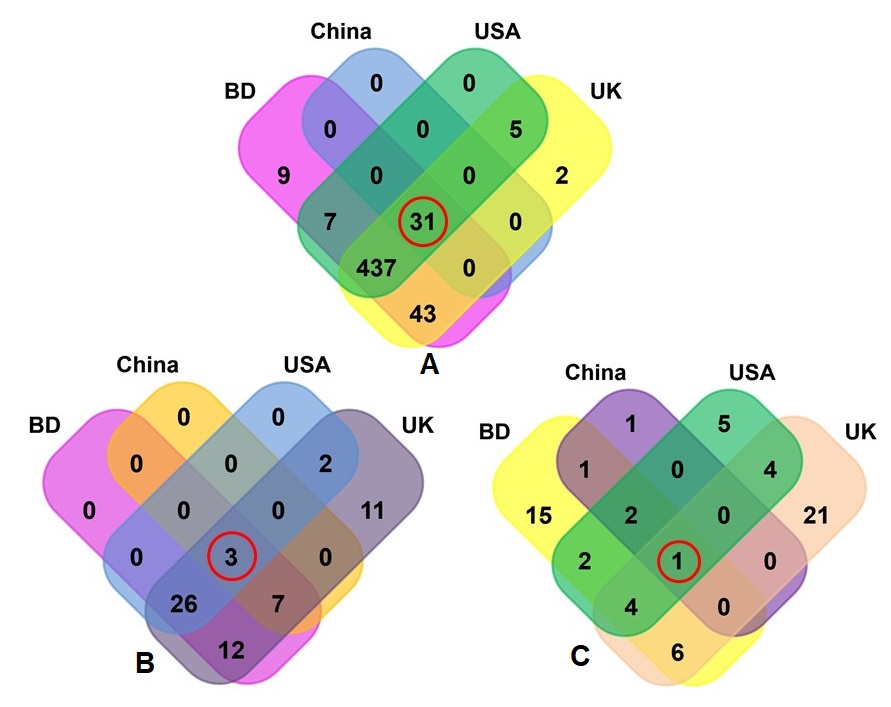
