## Supplementary material for "Diversity and genomic determinants of the microbiomes associated with COVID-19 and non-COVID respiratory diseases": Figure Legends

**Microbiome diversity of COVID 19 and other respiratory tract diseases, and genomic determinants**

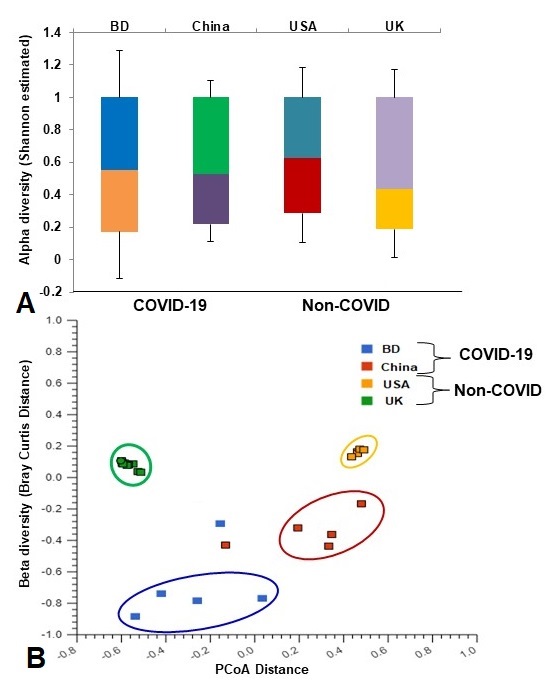

**Fig. 1. Differences in microbiome diversity and community structure in COVID-19 (BD and China), and non-COVID (UK and USA) disease metagenomes.** (A) Box plots showing significant differences (P=0.036, Kruskal-Wallis test) in Shannon estimated alpha diversity in four metagenomes. (B) Principal coordinates analysis (PCoA) measured on the Bray-Curtis distance method separated samples by microbial population structure. Each dot represents an individual, and colors indicate the populations in four metagenomes. Statistical analysis using Kruskal–Wallis tests showed significant microbial diversity variations across the four metagenomes (P=0.021, Kruskal-Wallis test).

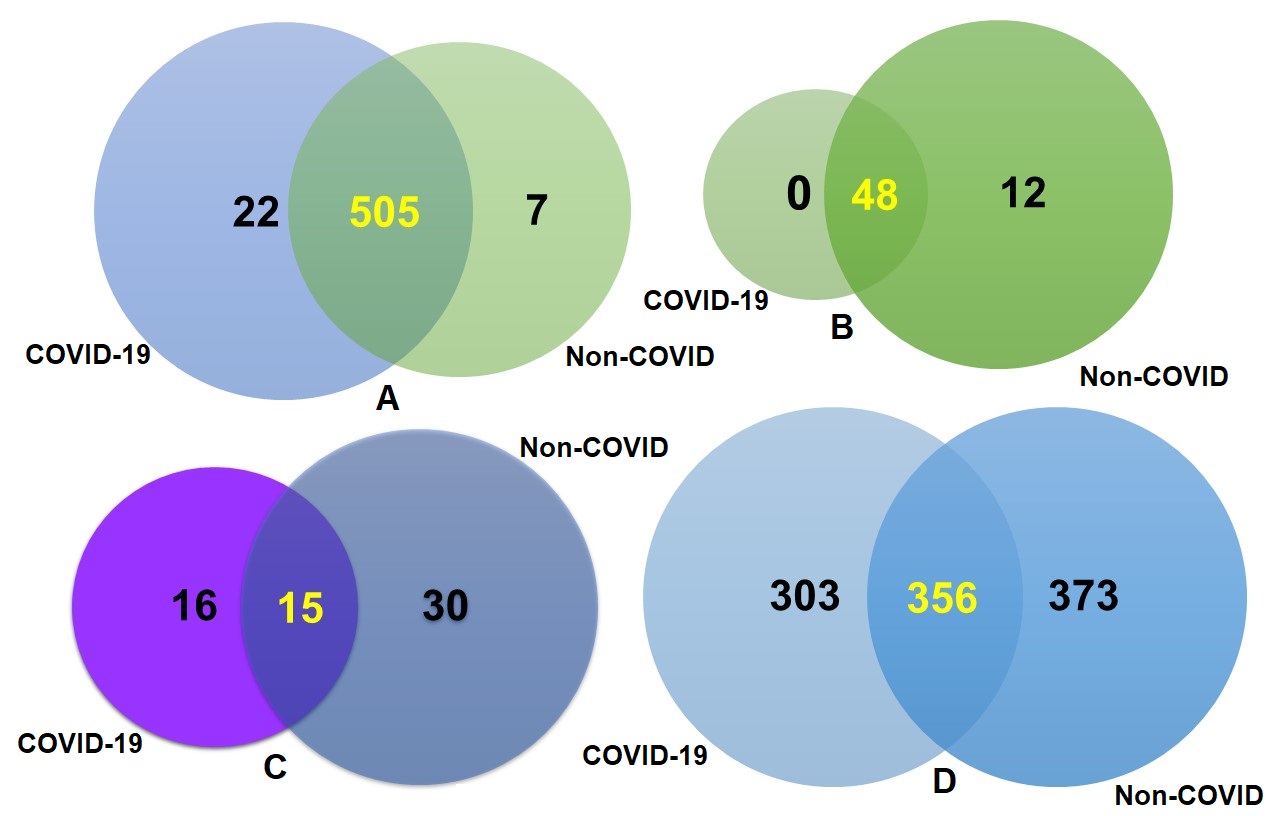

**Fig. 2. Taxonomic composition of COVID-19 (BD and China) and non-COVID (URTI; USA and COPD; UK) disease metagenomes.** Venn diagrams representing the core unique and shared microbiomes in COVID-19 and non-COVID diseases. (A) Venn diagram showing unique and shared bacterial genera. Out of 534 detected bacterial genera, only 13 genera had unique association with COVID-19 while rest of the 521 genera (highlighted in yellow) shared between the condition. (B) Venn diagram comparison of unique and shared archaeal genera where 48 (highlighted in yellow) genera shared between the condition, and 12 genera had unique association with non-COVID URTI and COPD) diseases. (C) Venn diagrams representing unique and shared viral genera identified in both metagenomes. Of the detected viral genera (n=61), 16 and 30 genera had unique association with COVID-19 and non-COVID diseases, respectively, and 15 genera (highlighted in yellow) were found to be shared between the study metagenomes. (D) Venn diagrams showing the unique and shared viral strains in COVID-19 and Non-COVID diseases. Out of viral strains detected, 356 strains (highlighted in yellow) shared between the condition while 303 and 373 strains had unique associations with COVID-19 and Non-COVID metagenomes, respectively. More information on the taxonomic results are available in Data S1.

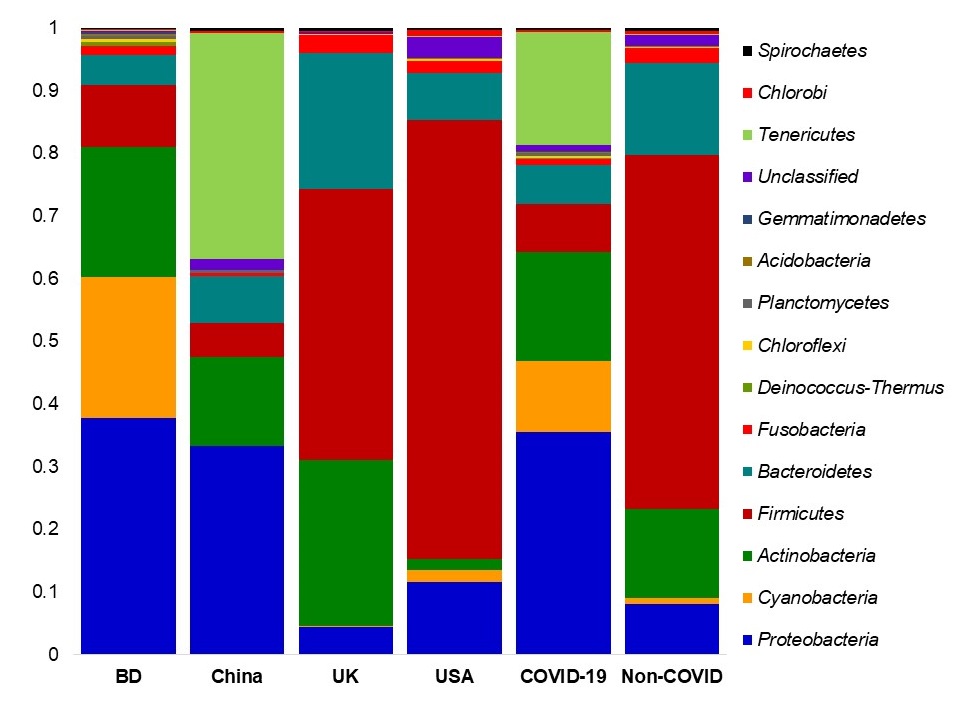

**Fig. 3**. **The phylum level taxonomic profile bacteria in in COVID-19 (BD and China), and respiratory tract (UK and USA) disease metagenomes.** Stacked bar plots showing the relative abundance and distribution of the 15 top abundant phyla, with ranks ordered from bottom to top by their increasing proportion. The first four bar plots represent the abundance of bacterial phyla in the corresponding category of COVID-19 (BD and China) and non-COVID (URTI; USA and COPD; UK) disease, and the last two bar plots depict overall relative abundance of phyla in COVID-19 and non-COVID metagenomes, respectively. The distribution and relative abundance of the bacterial phyla in the study metagenomes are also available in Data S1.

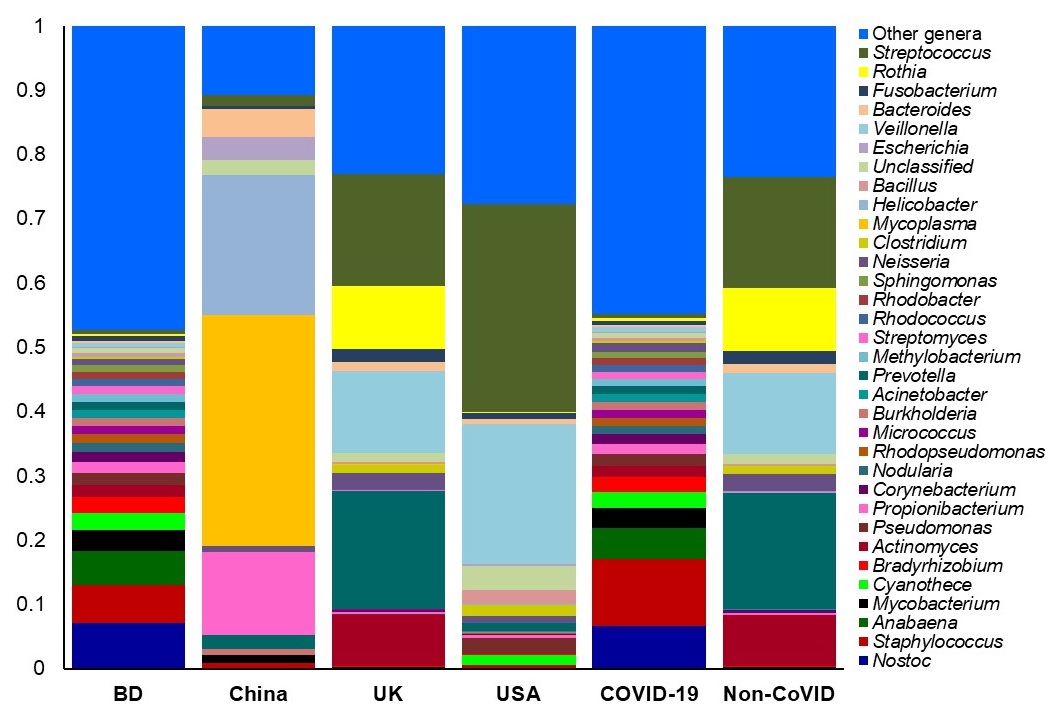

**Fig. 4**. **The genus level taxonomic profile bacteria in COVID-19 (BD and China) and non-COVID (URTI; USA and COPD; UK) disease metagenomes.** Stacked bar plots showing the relative abundance and distribution of the 34 most top abundant bacterial genera, with ranks ordered from bottom to top by their increasing proportion. The first four bar plots represent the abundance of bacteria in the corresponding category of COVID-19 (BD and China) and non-COVID (URTI; USA and COPD; UK) disease, and the last two bar plots depict overall relative abundance of bacterial genera in COVID-19 and non-COVID metagenomes, respectively. Only the 33 most abundant genera are shown in the legend, with the remaining genera grouped as ‘Other genera’. The distribution and relative abundance of the bacterial genera in the study metagenomes are also available in Data S1.

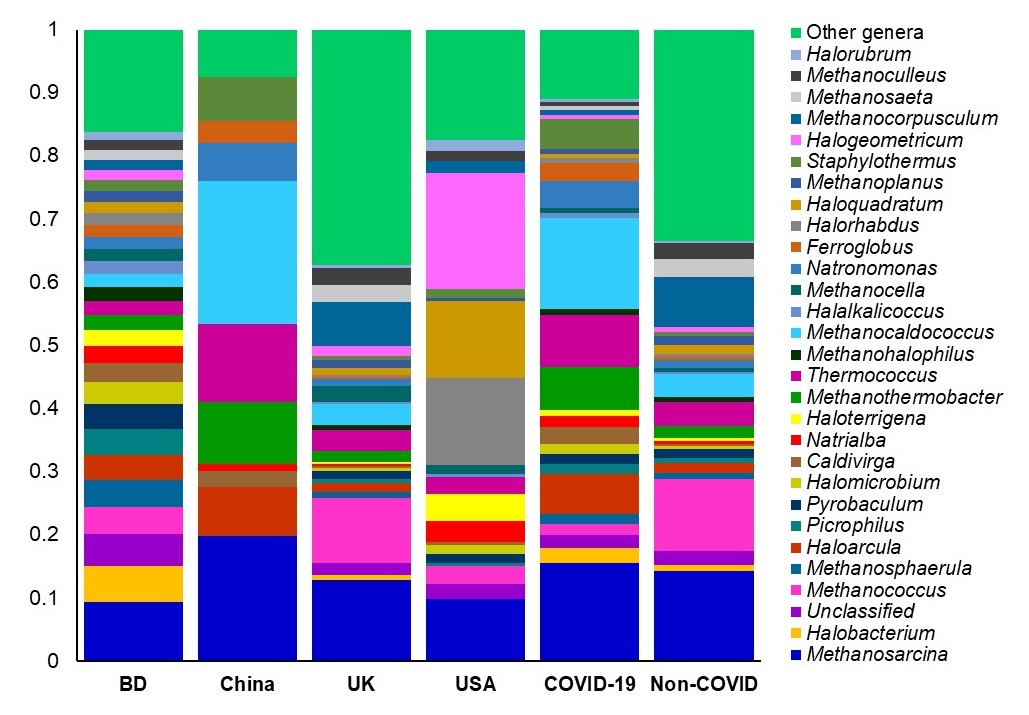

**Fig. 5**. **The genus level taxonomic profile archaea in COVID-19 (BD and China) and respiratory tract disease (UK and USA) metagenomes.** Stacked bar plots showing the relative abundance and distribution of the 30 most abundant genera, with ranks ordered from bottom to top by their increasing proportion. The first four bar plots represent the abundance of archaea in the corresponding category of COVID-19 (BD and China) and non-COVID (URTI; USA and COPD; UK) disease, and the last two bar plots depict overall relative abundance of archaeal genera in COVID-19 and non-COVID metagenomes, respectively. Only the 29 most abundant genera are shown in the legend, with the remaining genera grouped as ‘Other genera’. The distribution and relative abundance of the archaeal genera in the study metagenomes are also available in Data S1.

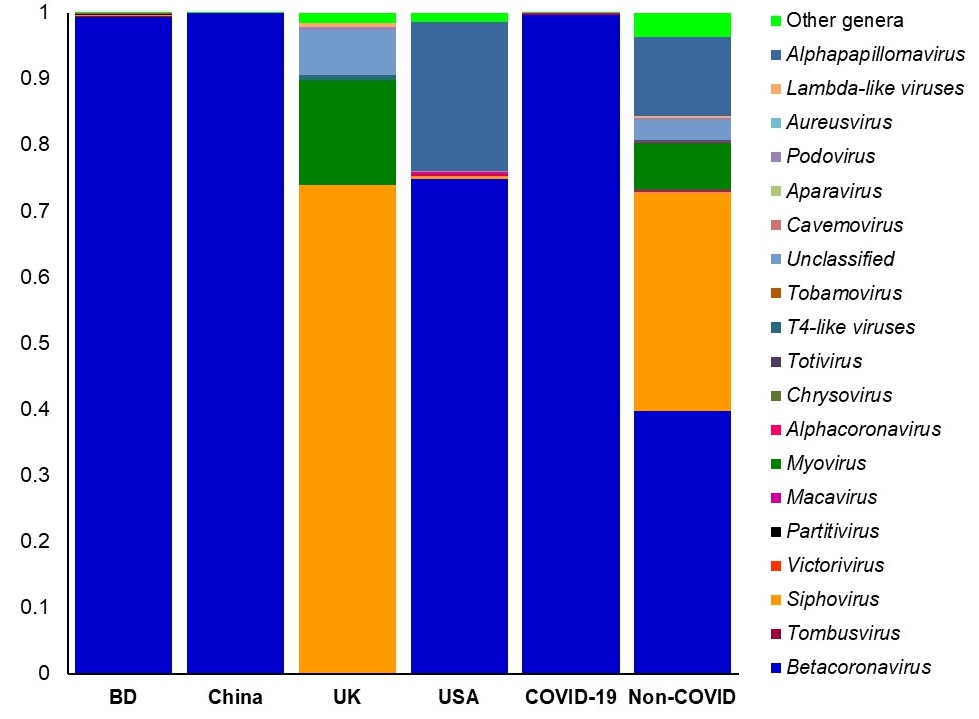

**Fig. 6**. **The taxonomic profile virus in in COVID-19 (BD and China) and respiratory tract disease (UK and USA) metagenomes.** Stacked bar plots showing the relative abundance and distribution of the 20 viral genera, with ranks ordered from bottom to top by their increasing proportion among the BD, China, UK and USA metagenomes. The first four bar plots represent the abundance of viruses in the corresponding category of COVID-19 (BD and China) and non-COVID (URTI; USA and COPD; UK) disease, and the last two bar plots depict overall relative abundance of viral genera in COVID-19 and non-COVID metagenomes, respectively. Only the 19 most abundant genera are shown in the legend, with the remaining genera grouped as ‘Other genera’. The distribution and relative abundance of the viral genera in the study metagenomes are also available in Data S1.

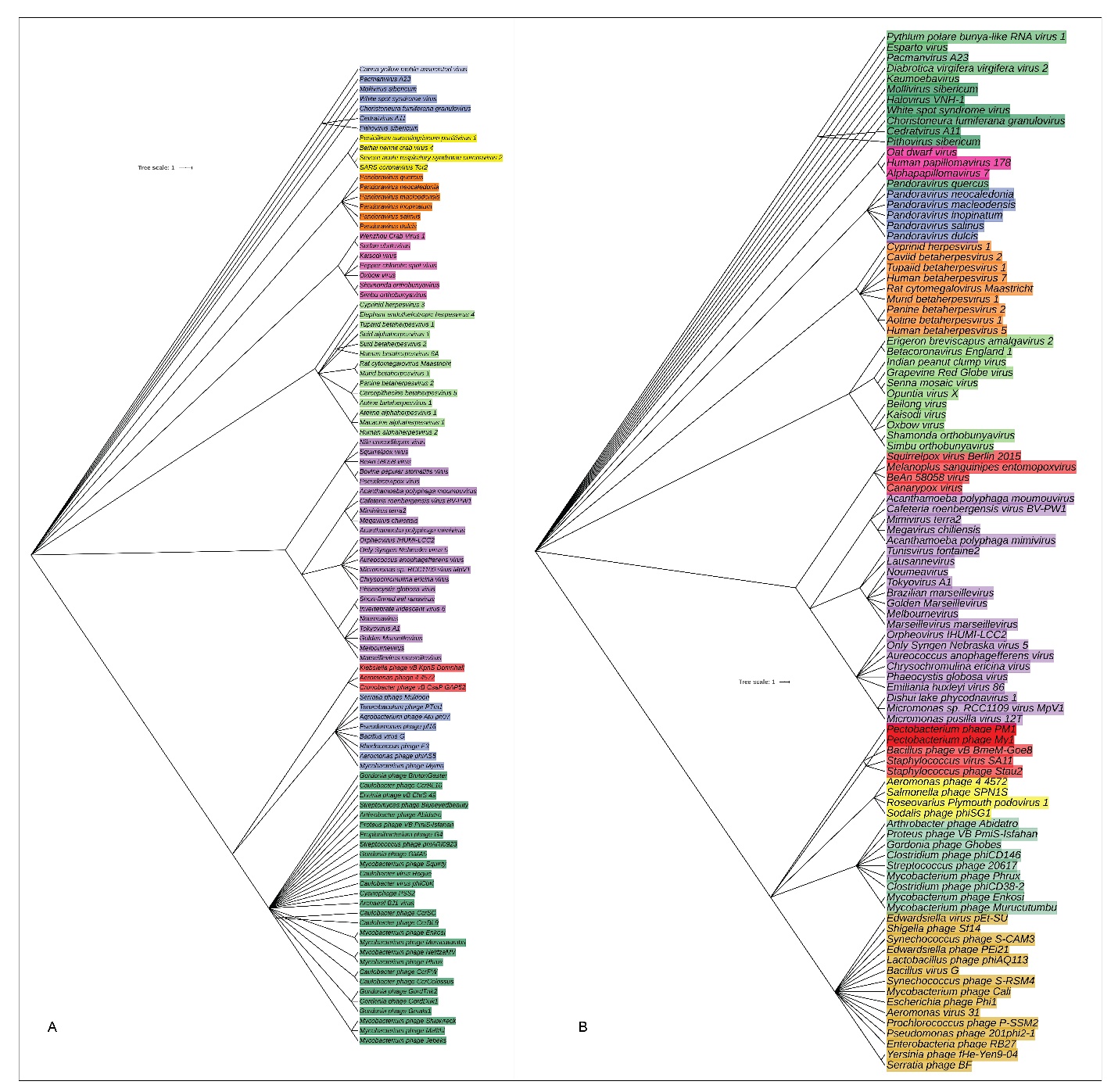

**Fig. 7:** The species and/or strain level taxonomic representation of viruses in COVID-19 and non-COVID (URTI and COPD) metagenomes. Sequences are assigned to different taxonomic index in PathoScope (PS) analysis using minimum identity of 95% and minimum alignment length 20 as cutoff parameters. The slanted phylogenetic trees were generated with the top 100 abundant strains of viruses in the COVID-19 (A) and non-COVID (B) metagenomes based on the maximum likelihood method using the NCBI taxonomy tree and visualized with iTOL (interactive Tree Of Life). The bootstrap considered 1000 replicates. The scale bar represents the expected number of substitutions averaged over all the analyzed sites. The length of the scale bar represents 1 nucleotide substitution per 100 positions. Different colors are assigned according to the taxonomic ranks of the viruses. The species and/or strains used in the phylogenetic tree are also available in Data S1.

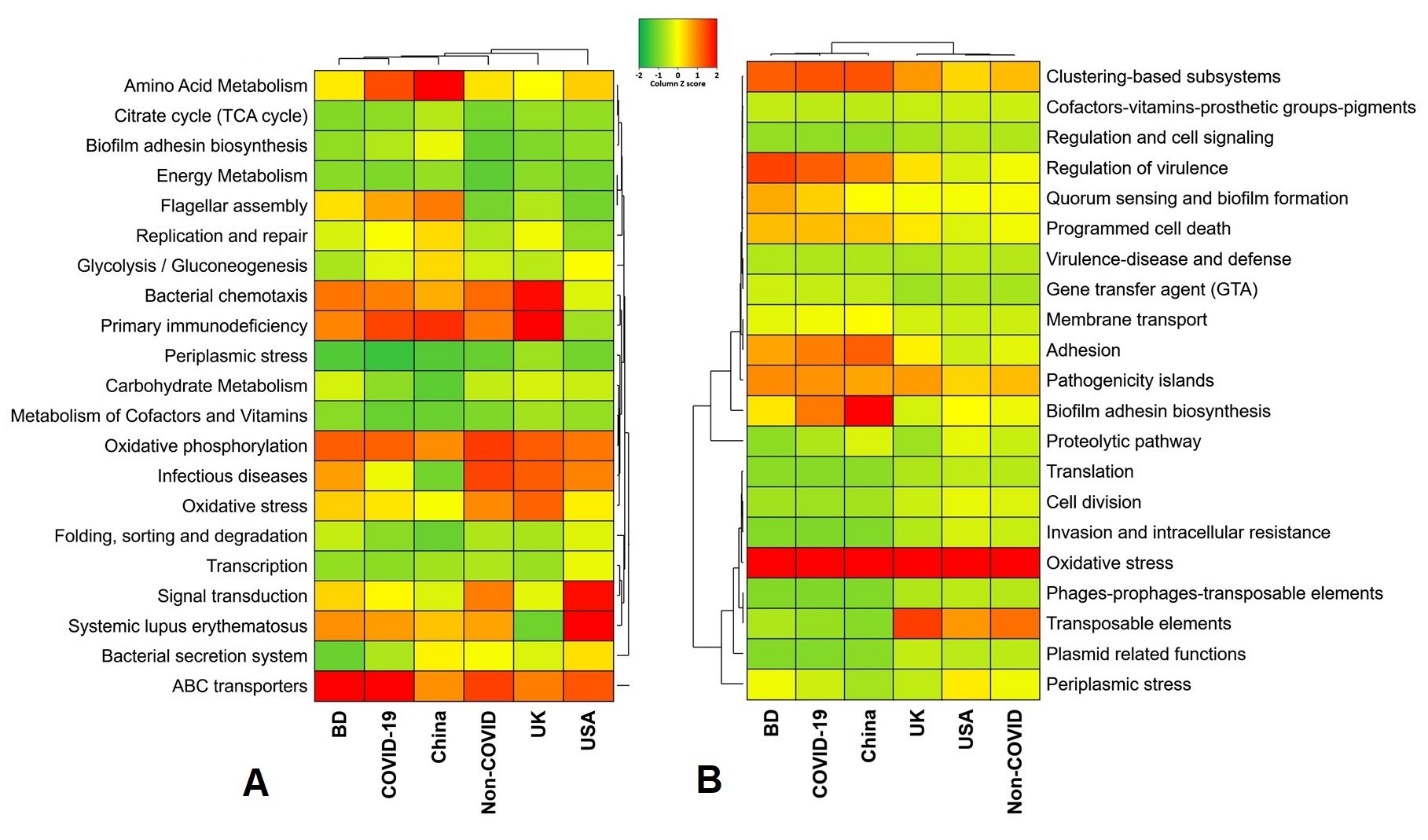

**Fig. 8.** **Functional annotation of the COVID-19 (BD and China) and respiratory tract disease (UK and USA) metagenomes**. (A) Heatmap representing the average relative abundance hierarchical clustering of the predicted KEGG Orthologs (KOs) functional pathways of the microbiome across four metagenome groups. (B) Heatmap showing the average relative abundance hierarchical clustering of the predicted SEED functions in different levels among the microbiomes of four metagenomes. The color bars (column Z score) at the top represent the relative abundance of putative genes. The color codes indicate the presence and completeness of each KEGG and SEED module, expressed as a value between -2 (lowest abundance) and 2 (highest abundance). The red color indicates the more abundant patterns, whilst green cells account for less abundant putative genes in that particular metagenome.

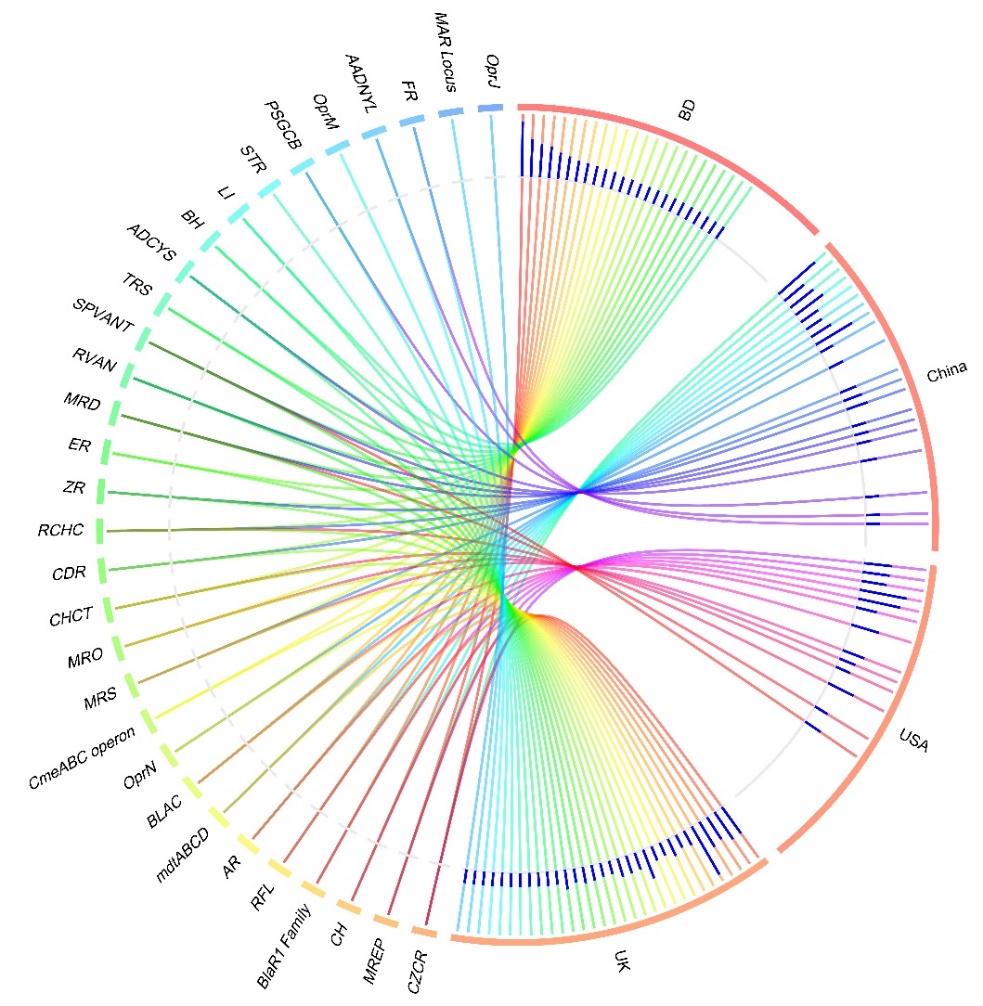

**Fig. 9.** **Distribution of the resistance to antibiotic and toxic compounds (RATC) genes in COVID-19 (BD and China) and respiratory tract disease (UK and USA) metagenomes.** The circular plot illustrates the diversity and relative abundance of the RATC genes detected among the microbiomes of the four metagenomes through SEED subsystems analysis. The association of the RATC genes according to metagenome is shown by different colored ribbons and the relative abundances these genes are represented by inner blue colored bars. Part of the RATC functional groups are shared among microbes of the four metagenomes (BD, China, UK and USA), and some are effectively undetected in the microbiomes of the other metagenomes. Abbreviations- *CZCR*: cobalt-zinc-cadmium resistance, *MREP*: multidrug resistance to efflux pumps, *CH*: copper homeostasis; *BlaR1* Family: BlaR1 family regulatory sensor-transducer disambiguation, *RFL*: resistance to fluoroquinolones, *AR*: arsenic resistance, *mdtABCD*: the mdtABCD multidrug resistance cluster, *BLAC*: beta-lactamase resistance, *OprN*: mexe-mexf-oprn multidrug efflux system, *CmeABC* operon: multidrug efflux pump in *Campylobacter* *jejuni*, *MRS*: methicillin resistance in *Staphylococci*, *MRO*: mercury resistance to operon, *CHCT*: copper homeostasis: copper tolerance, *CDR*: cadmium resistance, *RCHC*: resistance to chromium compounds, *ZR*: zinc resistance, *ER*: erythromycin resistance, *MRD*: mercuric reductase, *RVAN*: resistance to vancomycin, *SPVANT*: *Streptococcus pneumonia* vancomycin tolerance locus, *TRS*: teicoplanin-resistance in *Staphylococcus*, *ADCYS*: adaptation to d-cysteine, *BH*: bile hydrolysis, *LI*: lysozyme inhibitors, *PSGCB*: polymyxin synthetase gene cluster in *Bacillus*, *OprM*: mexA-mexB-oprm multidrug efflux system, *AADNYL*: aminoglycoside adenylyltransferases, *FR*: fosfomycin resistance, *MAR* Locus: multiple antibiotic resistance to locus, *OprJ*: mexC-mexD-OprJ-multidrug-efflux-system.
