## Supplementary Figure Legends for "Diversity and genomic determinants of the microbiomes associated with COVID-19 and non-COVID respiratory diseases"

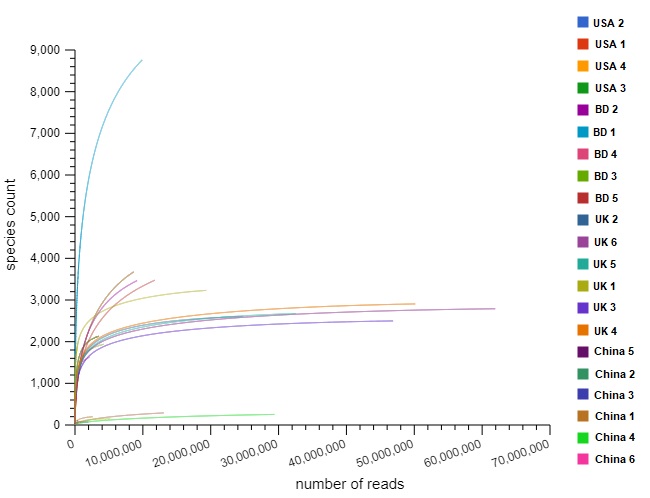

**Fig. S1. Differences in species richness in COVID-19 (BD and China), and non-COVID (UK and USA) disease metagenomes.** Rarefaction curves showing the influence of sequencing depth (number of reads per sample, X axis) on species richness (Y axis) in COVID-19 and non-COVID samples. The rarefaction curves representing the number of species per sample indicated that the sequencing depth was sufficient enough to fully capture the microbial diversity as existed.

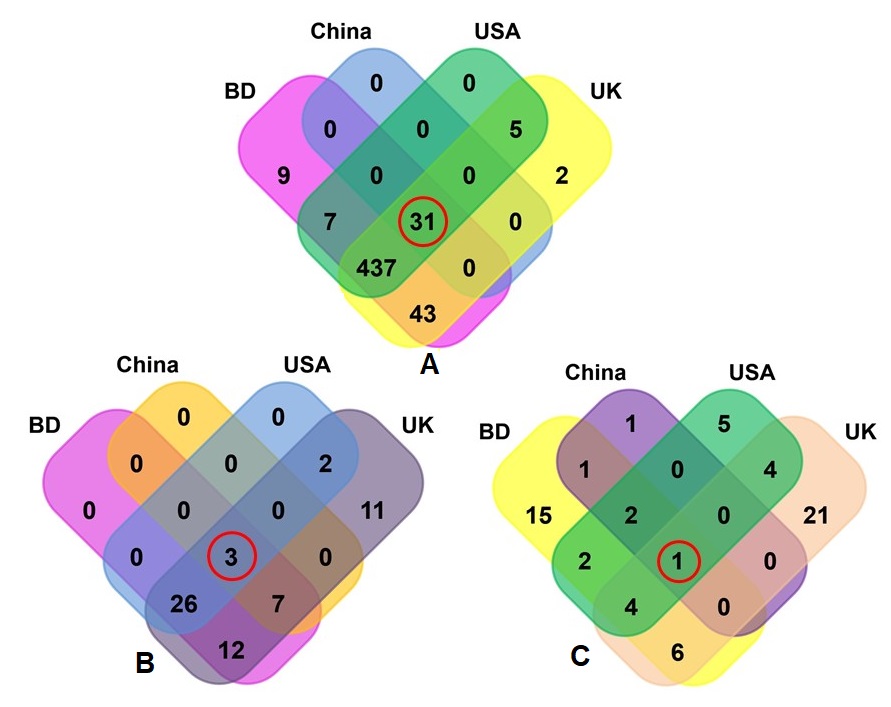

**Fig. S2. Taxonomic composition of COVID-19 (BD and China) and respiratory tract disease (UK and USA) metagenomes.** Venn diagrams representing the core unique and shared microbiomes in BD, China, UK and USA metagenomes. (A) Venn diagram showing unique and shared bacterial genera; (B) Venn diagram comparison of unique and shared archaeal genera; and (C) Venn diagrams representing unique and shared viral genera identified in four metagenomes through MG-RAST analysis. Microbiome sharing between the conditions are indicated by red circles.
